# Genome-resolved metagenomics reveals microbial potential for CO_2_ and CH_4_ emissions in prawn ponds

**DOI:** 10.64898/2026.08.18.745432

**Authors:** Abul Bashar, Amaru Miranda Djurhuus, Patrick Denis Browne, M.M.R. Jahangir, Niels O. G. Jørgensen, Mohammad Mahfujul Haque, Lars Hestbjerg Hansen

## Abstract

Recognizing the central role of microorganisms in greenhouse gas (GHG) cycling in aquaculture systems, we provide a genome- and gene-centric perspective on the metabolic potential for CO₂ and CH₄ cycling in prawn aquaculture ponds across seasons and contrasting culture practices. Using TaxVAMB, we recovered 78 high- and medium-quality metagenome-assembled genomes (MAGs), including previously underappreciated taxa such as *Bathyarchaeia* and *Terriglobia*. Metabolic profiling revealed that CO₂ and CH₄ cycling constitute a minor fraction of the pond’s metabolic potential, dominated instead by heterotrophic processes such as fermentation, oxygen metabolism, and iron reduction. The relative metabolic weight of these carbon-cycling pathways was lower than that reported for permafrost, wetland, peatland, deep-sea, and human gut microbiomes. An integrated metabolic network suggested that genetic potential for CO₂ production is primarily driven by pyruvate and acetyl-CoA oxidation, while methanogenesis and methane oxidation genes together encode the potential for internal carbon-recycling loops via canonical archaeal and bacterial pathways. Seasonal dynamics, rather than management treatment, strongly influenced functional gene abundances, with CO₂ fixation and CH_4_ oxidation genes increasing toward the late season. *Bathyarchaeia* emerged as the most versatile taxon for CO₂ cycling and methanogenesis, with stable relative abundance across seasons and treatments. This study underscores the role of seasonally evolving microbial networks in regulating carbon turnover and the potential for CO_2_ and CH_4_ emissions in prawn aquaculture ponds.

## INTRODUCTION

Aquaculture ponds are highly dynamic natural biogeochemical reactors characterized by intensive organic matter loading, rapid nutrient turnover, and steeply fluctuating redox conditions. These features promote complex, changing microbiomes capable of performing a range of metabolic pathways, including organic matter degradation, carbon fixation, methanogenesis, methane oxidation, nitrification, and denitrification^1–5^, thereby critically determining net in-pond GHG emissions and affecting the environmental impact of aquaculture. In prawn aquaculture, the heavy reliance on formulated feeds and the accumulation of uneaten feed and metabolic waste substantially enhance organic substrate availability, accelerating microbial-mediated biogeochemical cycles^6,7^. As a result, prawn ponds act as localized hotspots of GHG emissions^8^, with our recent life-cycle assessment estimating in-pond emissions of approximately 700-1300 g CO_2_-equivalent (CO₂e) per kilogram of harvested biomass, depending on culture practices^9^.

Bacteria and archaea are the primary microbial drivers of CO_2_ and CH_4_ cycling in aquatic ecosystems^7,10^, with microbial methanogenesis accounting for an estimated 69% of global methane emissions^11^. In aquaculture ponds, methyl co-enzyme M reductase (*mcr)*-containing methanogenic archaea produce methane under anaerobic conditions via hydrogenotrophic and acetoclastic pathways^3,12–14^. At the same time, oxic–anoxic interfaces at the sediment-water boundary provide niches for aerobic and microaerophilic methanotrophs, which oxidize a substantial fraction of produced methane and act as biological methane sinks^15^. In parallel, chemolithoautotrophic microorganisms contribute to CO₂ fixation^16,17^, influencing net carbon fluxes in pond ecosystems. However, the magnitude and direction of these microbial processes are highly sensitive to environmental conditions, including organic substrate availability, electron acceptor concentrations, trace element supply, and oxygen dynamics^18,19^. For example, elevated sulfate concentrations can favor sulfate-reducing bacteria (SRB) that outcompete methanogens for shared carbon substrates and form syntrophic associations with anaerobic methanotrophs, including ANME-1 and ANME-3 lineages, thereby reducing methane production^20–23^. Therefore recognizing the crucial role of microorganisms in GHG emissions, understanding their compositional and metabolic profiles under varying environmental conditions is crucial to enhance our understanding and inform efforts to mitigate microbial-mediated emissions in prawn aquaculture ponds.

Recent metagenomic and metatranscriptomic reports have advanced understanding of CO₂- and CH₄-cycling microorganisms across diverse ecosystems, including hot springs^24^, permafrost soils^25^, deep-sea sediments^26^, coastal wetlands and mangroves^16,27–29^, and agricultural and aquaculture systems^30–36^. In aquaculture environments, most existing investigations emphasized taxonomic composition, pathogen dynamics^34,36,37^, or antimicrobial resistance^30–32^, primarily in shrimp and finfish systems, with little attention to functional potential related to carbon cycling. Our previous work explored CO_2_ and CH_4_ cycling microbial populations in prawn ponds but was limited to taxonomic profiling based on 16S rRNA gene amplicon sequencing, precluding functional inference of emission pathways^7^. Moreover, it was temporally constrained, providing an incomplete picture of how microbial communities and metabolic pathways respond to dynamic pond conditions. In fact, microbial community structure and function in prawn ponds are expected to vary substantially over production cycles due to progressive organic matter accumulation, seasonal climatic variability, and declining oxygen availability associated with increasing biological oxygen demand. Such physicochemical dynamics are likely to generate pronounced spatial and temporal heterogeneity in microbial assemblages and metabolic pathways. For example, growing organic loads and oxygen depletion may promote the expansion of anaerobic micro-niches over time, enhancing methanogenic activity in pond sediments^38^.

Farm management interventions may reduce the availability of microbial substrate by reducing organic matter in ponds. One such promising technique is the Integrated multi-trophic aquaculture (IMTA) system, which combines fed species with organic and inorganic extractive organisms^9^. In IMTA system, bottom-feeding extractive organisms (e.g. snails, bivalves) consume excess organic matter and uneaten feed, while inorganic extractives (e.g. water spinach, seaweeds) assimilate dissolved inorganic nutrients^9,39^, collectively reducing sediment organic loading and nutrient availability for anaerobic microbial processes. By enhancing oxygen availability through photosynthesis and limiting substrates for methanogenesis, IMTA has the potential to suppress CO₂- and CH₄-producing pathways and shift microbial community structure^40^ toward lower greenhouse gas emissions. However, whether and how CO₂- and CH₄-cycling microbial populations and pathways differ between IMTA and conventional prawn farming systems remains unexplored.

To manage prawn aquaculture ponds in a climate-responsible manner, it is essential to identify the dominant microbial metabolic pathways that regulate GHG emissions and to understand how these pathways change over the course of a production cycle. Equally important is resolving how the abundance of key functional genes and the composition of microbial populations shift over time and across contrasting culture practices. An ideal experimental setting to address these questions is the subtropical, DANIDA-funded ECOPRAWN experimental facility established in southwest Bangladesh (22°47′07″ N, 89°27′50″ E), which comprises replicated ponds operated under contrasting culture gradients and is embedded within an ongoing life cycle–based effort to quantify and mitigate in-pond GHG emissions through the implementation of an IMTA system^9^. Here, we applied shotgun metagenomic sequencing to resolve temporal dynamics in microbial community composition and functional potential, with a particular focus on CO₂ and CH₄ cycling microorganisms, across contrasting culture practices over a complete production cycle. We focused on pond sediments, where most organic matter accumulates and persists, providing the primary substrate for microbial metabolism. Using complementary genome- and gene-centric approaches, we reconstructed microbial metabolic pathways associated with CO₂ and CH₄ cycling by recovering metagenome-assembled genomes (MAGs) and quantifying temporal changes in the abundance of functional genes to infer gains and losses in metabolic potential. Through these analyses, we tested two hypotheses: (1) microbial metabolic processes and CO₂ and CH₄ emission potentials are governed by time and culture practice; and (2) The relative abundance of functional genes participating in CO_2_ and CH_4_ cycling and emissions varies over time and across culture systems, in response to physicochemical conditions and organic loading, with increasing taxonomic diversity of gene-harboring microorganisms toward the end of the production cycle.

## METHODS

### Experimental background, sediment sample collection, and DNA extraction

The field experiment was conducted in 2024 at the ECOPRAWN experimental facility, where eighteen earthen ponds were constructed, comprising six treatments^9^. The experimental platform was designed to support a life cycle–based assessment of in-pond GHG emissions, covering a combined water area of 1.29 ha with a mean water depth of 1.83 m. The treatments were designed to progressively disentangle biotic effects and evaluate the role of organic and inorganic extractive components within an IMTA framework (Table 1). Treatment T1 reflects conventional polyculture practice with prawn and carp, whereas treatment T2 contained no cultured organisms to characterize background pond biogeochemistry and microbial communities in the absence of aquaculture activities. T3 introduced local snails (*Bellamya bengalensis*) as organic extractives that graze on uneaten feed and particulate organic matter, while T4 incorporated water spinach (*Ipomoea aquatica*) as an inorganic extractive component that assimilates nitrogenous compounds from the water column. Treatment T5 represented a prawn monoculture system stocked solely with prawns. An IMTA system combining prawn and carp as main fed species, water spinach as an inorganic extractive component, and snails as an organic extractive species was designated treatment T6. Across all aquaculture treatments, prawns were fed a commercial formulated diet at rates declining from approximately 7% to 3% of standing biomass per day. Water spinach was harvested weekly to stimulate vegetative growth and maintain approximately 15-20% surface coverage of the pond.

**Table 1.** Experimental treatments indicating culture system type, inclusion of components, including main fed components and extractive components, used in the ECOPRAWN experiment.

| Treatments | Culture systems | Components |
| --- | --- | --- |
| T1 | Improved-extensive polyculture | Prawn and Indian major carps |
| T2 | Non-aquaculture | None |
| T3 | Improved-extensive polyculture | Prawn, Indian major carps, and snail |
| T4 | Improved-extensive polyculture | Prawn, Indian major carps, and water spinach |
| T5 | Improved-extensive monoculture | Prawn |
| T6 | Integrated Multi-trophic Aquaculture | Prawn, Indian major carps, snail, and water spinach |

Six coordinated sampling campaigns (S1: day 30; S2: day 60; S3: day 90; S4: day 120; S5: day 150; and S6: day 180) were conducted. During each campaign, in-pond GHG fluxes were measured, water quality parameters were monitored, and biological yields of prawn and carp were assessed. In parallel with gas sampling, sediment samples for microbial analyses were collected aseptically from each pond using a sterile stainless-steel auger^7^. Because oxygen penetration in organic-rich aquaculture sediments is typically limited to the upper few millimetres, we collected sediment cores from the top 2 cm to capture both aerobic and anaerobic microbial communities. Genomic DNA was extracted from 0.3 g of homogenized sediment using the DNeasy PowerSoil Kit (Qiagen, Hilden, Germany) and if necessary, was further purified using the DNA Clean & Concentrator kit (Zymo Research, USA). To minimize small-scale spatial heterogeneity within treatments, DNA extracts from triplicate ponds were pooled, and final DNA concentration and purity were assessed using a Qubit fluorometer (Thermo Fisher Scientific) and a NanoDrop spectrophotometer, respectively.

### Hackflex library preparation and metagenomic sequencing

For shotgun metagenomic sequencing, libraries were prepared using an adapted Hackflex protocol, a low-cost modification of the Nextera DNA Flex (Illumina) library preparation, using laboratory-made reagents and custom index primers^41,42^. The protocol followed previously described Hackflex workflows^41^ with minor optimizations for sediment-derived metagenomic DNA (Figure S2). Briefly, 5-10 ng of high-quality genomic DNA per sample was used as input. Tagmentation was performed using bead-linked transposomes (BLT; Illumina DNA Library Prep) diluted 1:50 in nuclease-free water and combined with a laboratory-made 2x tagmentation buffer containing Tris-HCl, MgCl₂, and dimethylformamide (see Supplementary note 2 for detailed recipe and procedure). Reactions were incubated at 55 °C for 15 min in a thermocycler with a heated lid, followed by immediate reaction termination using tagmentation stop buffer (0.2% sodium dodecyl sulfate) at 37 °C for 15 min. PCR amplification was conducted directly on BLT beads using a high-fidelity DNA polymerase (PCRBio HiFi), replacing the Illumina-recommended polymerase to optimize amplification efficiency and fidelity for sediment DNA. Each PCR reaction contained 10 µl of 5× PCRBio HiFi buffer, 0.50 µl of PCRBio HiFi polymerase, and 29.50 µl of nuclease-free water. Illumina-compatible custom dual-index i5 and i7 barcodes (v1 set) developed by Gao et al^41^ were added at 5 µl each to a final primer concentration of 1 µM per index. PCR amplification was performed in a thermocycler with a heated lid using the following thermal conditions: an initial denaturation at 98 °C for 3 min, followed by 12 amplification cycles consisting of denaturation at 98 °C for 45 s, annealing at 62 °C for 30 s, and extension at 68 °C for 2 min^41^. The number of PCR cycles was optimized based on the input DNA concentration in accordance with Illumina DNA Prep guidelines to minimize amplification bias.

Amplified libraries were purified, and size selection was performed using AMPure XP paramagnetic beads (Beckman Coulter) through a double size selection procedure designed to remove both short (including primer dimers) and larger DNA fragments^41^. Briefly, an equal volume of nuclease-free water was added to the PCR products, followed by the addition of AMPure XP beads at a 1.0x bead-to-PCR volume ratio to bind larger DNA fragments. The supernatant containing small and medium-sized fragments was then subjected to a second size-selection step by adding AMPure XP beads at 0.33x bead-to-PCR volume, which selectively binds medium-sized DNA fragments. Bead-bound DNA was washed twice with freshly prepared 70% ethanol, air-dried immediately, and eluted in 20 µl of nuclease-free water. Final library concentrations were quantified using the Qubit High Sensitivity dsDNA assay (Thermo Fisher Scientific). Equimolar amounts of individual libraries were pooled and subjected to shotgun metagenomic sequencing (PE 150) on an Illumina NovaSeq platform (Illumina Inc., San Diego, CA, USA) at Novogene, UK.

### Metagenomic assembly, TaxVAMB binning, and functional annotation

Raw reads were quality-filtered and trimmed using fastp v0.23.2, removing low-quality bases (Phred score< 20) and reads with >30% of low-quality bases and shorter than 50 bp. High-quality reads were assembled into contigs using MEGAHIT v1.2.9 with default k-mers^43^ and the quality of the assembled contigs was assessed using QUAST v5.2.0^44^. The resultant high-quality contigs were then fed into the TaxVAMB pipeline for semi-supervised binning^45^. This is a deep learning-based binning tool that integrates intrinsic contig features (co-abundance profiles) with annotation features such as taxonomic labels. TaxVAMB employs a bi-modal variational autoencoder to generate embedded representations of contigs, followed by iterative clustering, producing high-quality MAGs that recover greater taxonomic diversity than traditional binning approaches^45^. This approach is particularly practical for datasets with fewer than 100 samples and incomplete genomes from complex environments. Concatenated contigs were then mapped to raw reads using StrobeAlign v0.7.5^46^ and were classified taxonomically using MMseqs2 taxonomy module^47^ with the GTDB v2.1.1 database to generate abundance estimates and taxonomic labels. MAGs were recovered and integrated with contig coverage and taxonomic information. The quality (completeness and contamination) of bins was assessed using CheckM2^48^, and high- and medium-quality (HMQ) MAGs were dereplicated using dRep v3.5.2^49^ with thresholds of ≥75% completeness and ≤5% contamination and assigned taxonomies by linking to the contig labels.

To infer the metabolic potential of MAGs, ORFs were predicted for all genomes using the Prodigal algorithm implemented within METABOLIC v4.0 and annotated by searching against multiple hidden Markov model (HMM) databases, including KEGG, KOfam, Pfam, TIGRFAM, MEROPs, with default parameters^50^. To quantify the dominance of individual metabolic pathways in prawn ponds, the metabolic weight (MW) score was calculated for each pathway by integrating related gene presence-absence information with genome coverage derived from read-mapping results^51^. The metabolic weight score for a specific metabolic function *m* was calculated with the following equation^51^:

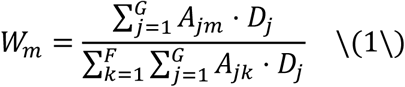

where *W_m_*denotes the weighted contribution of metabolic function *m*, *G* represents the total number of genomes, and *F* is the total number of annotated functions. The variable *D_j_* corresponds to the sequencing coverage of genome *j*, while *A_jm_* is a binary indicator describing the presence (1) or absence (0) of function *m* in genome *j*.

To assess the relative contribution of each microbial class to the weighted metabolic potential of a given pathway, we calculated the proportional contribution *P_m_*_,*p*_of class *p* to metabolic function *m* as follows^51^:

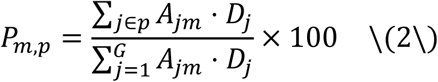

where genomes belonging to class *p* are represented by the subset *j* ∈ *p*. This calculation quantifies the percentage contribution of each taxonomic group to the overall metabolic weight of a given pathway.

ORFs were also predicted at the bin level using Prodigal v2.6.3^52^ in metagenomic mode and predicted proteins were functionally annotated using KofamScan v1.3.0^53^, assigning KEGG Orthology identifiers to each gene for gene-level downstream analysis.

### Statistical Analysis

Metagenomic datasets, including functional gene abundances, MW score, and metabolic contribution of MAGs, were imported into R v4.4.3 for downstream analysis. Heatmaps representing MW scores and percentage contributions were generated using *ComplexHeatmap*, with color gradients indicating the scores and percentages. Mean abundances of functional gene groups were visualized as barplots with overlaid individual sample points using *ggplot2*. Gene abundances were z-score normalized per gene, and dimensionality reduction was performed using *UMAP*, followed by k-means clustering to identify sample clusters, which were visualized with *ggplot2*. For key CO_2_ and CH_4_ cycling genes, abundances were aggregated by season and treatment, and functional ratios (production to fixation/oxidation) were calculated. Mean ratios were visualized using line plots with error ribbons generated in *ggplot2*. Gene–function–taxon relationships were summarized and exported for visualization as a Sankey diagram using *SankeyMATIC*. Alpha diversity indices were calculated using *vegan* and differences across seasons and treatments were assessed using the Kruskal-Wallis (KW) test, followed by Dunn’s post hoc comparison. Beta diversity was evaluated using Bray-Curtis dissimilarities, and non-metric multidimensional scaling (NMDS) was performed with two dimensions, using 100 iterations. Relative abundances of core microbial classes were normalized per sample and aggregated by season and treatment for visualization in *ggplot2*.

## RESULTS

### Diversity and abundance of high- and medium-quality MAGs recovered from prawn ponds

Taxvamb binning recovered 82,181 bacterial and archaeal bins spanning a wide range of completeness and contamination levels (Fig. 1A and Supplementary note 3). From these, 78 HMQ MAGs that met stringent quality thresholds (>75% completeness and <5% contamination) were retained for metabolic investigations. These included 63 bacterial and 15 archaeal MAGs, with taxonomic classification using GTDB-Tk2 resolving 41 MAGs to the species, 34 to the genus, two to the family, and one to the order level. Among bacterial MAGs, Bacteroidia (n=27), Anaerolineae, Campylobacteria, and Gammaproteobacteria (n=6) were taxonomically diverse classes, whereas Methanosarcina (n=5), and Thermoplasmata (n=4) classes dominated Archaeal HMQ MAGs with additional representation from Methanobacteria, Nitrososphaeria, Hermodarchaeia, and Micrarchaeia (Fig. 1B). Anaerolineae (0.20–0.25) and Terriglobia (0.11–0.20) dominated bacterial abundance relatively, followed by Ignavibacteria (0.06–0.11), Bacteroidia (0.04–0.09), and Gammaproteobacteria (0.04–0.13). Species-level classification identified dominant taxa including *Ginsengibacter sp035765495*, *Palsa-295 sp035576265*, and *Planktothricoides raciborskii*. Archaeal classes were comparatively rare, with Methanosarcina and Thermoplasmata exhibiting low relative abundances (<0.03). Seasonal trends revealed a significant proliferation of sulfate-reducing MAGs within the Desulfobacteria and DSM lineages (KW, p < 0.05) (Fig. 1C), likely reflecting the accumulation of organic matter in the water. In contrast, treatment-wise significant dynamics showed Gammaproteobacteria peaking in T6 and Desulfobacteria reduced in T2 (see Extended Data ST1 for MAG-specific trends).

**Fig. 1:**
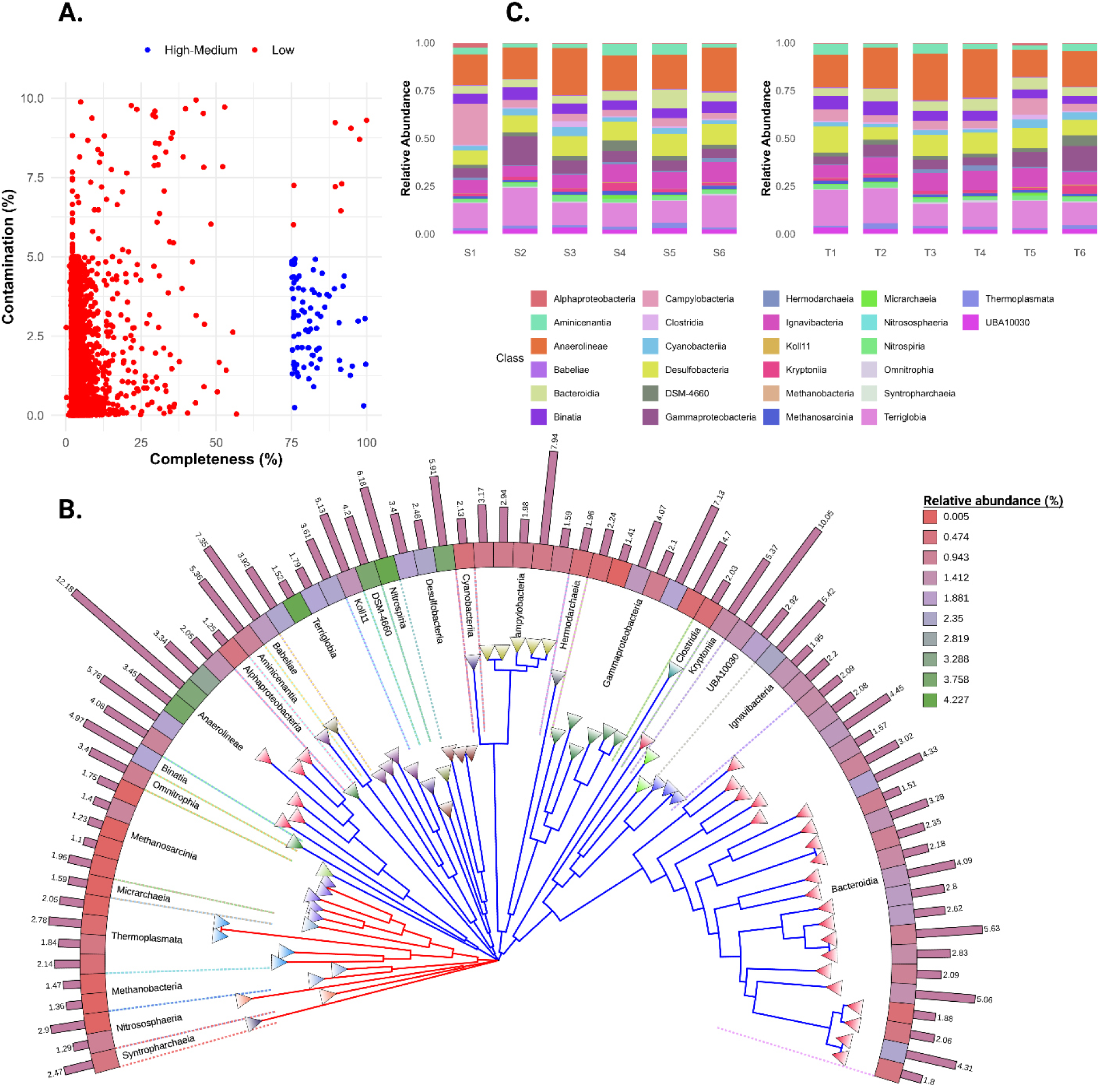
Quality metrics taxonomic diversity, phylogenetic relationships, and relative abundance of metagenome-assembled genomes (MAGs) recovered from prawn ponds. Quality assessment of recovered MAGs based on estimated completeness and contamination (A). High- and medium-quality (HMQ) MAGs, defined as >75% completeness and <5% contamination, are highlighted in blue. Maximum-likelihood phylogenetic tree of bacterial and archaeal MAGs reconstructed from concatenated conserved marker genes using FastTree (B). Branches corresponding to Archaea and Bacteria are shown in red and blue, respectively. Node colors in a gradient scale indicate class-level taxonomic assignment. The outer heatmap ring represents the relative abundance of individual MAGs, and bars indicate estimated genome size in Mbp. Relative abundance of MAG classes across treatment and sampling seasons (C).

### CO₂ and CH₄ cycling metabolisms constitute a minor fraction of community functions in prawn ponds

To assess the contribution of CO₂ and CH₄ cycling processes to the overall metabolic potential of prawn ponds, we calculated community-level metabolic weight scores across 26 metabolic pathways. Methanotrophy (1.24%) and methanogenesis (0.15%) represented only a minor fraction of total community metabolism, particularly when compared with dominant pathways such as fermentation (12.14%), oxygen metabolism (12.18%), and iron reduction (11.65%) (Fig. 2A). Similarly, carbon fixation (1.56%) and C1 metabolism (2.26%) were restricted to a small subset of MAGs (14 and 12 MAGs, respectively), suggesting that autotrophic carbon assimilation and methane cycling play relatively minor roles in prawn pond metabolic networks. Analysis of taxonomic contributions revealed that methanotrophy was primarily associated with Gammaproteobacteria (40.62%), Alphaproteobacteria (20.20%), and Thermoplasmata (30.06%), whereas methanogenesis was predominantly driven by Methanosarcina (86.26%) and Methanobacteria (4.00%) (Extended Data ST2). In contrast, carbon fixation was distributed across diverse taxa, including Anaerolineae (30.58%), Cyanobacteria (16.16%), and Nitrospiria (13.24%), whereas C1 metabolism was primarily represented by Binatia (27.36%), Terriglobia (40.96%), and Methanosarcina (7.09%). Given the intense metabolic coupling between sulfur and methane cycling in anaerobic environments, we also examined sulfur/sulfide cycling metabolism. Desulfobacteria and DSM-4660 exclusively performed sulfur/sulfide reduction (0.85%, 3 MAGs). Conversely, oxidation of sulfur/sulfide (8.18%, 38 MAGs) was more widespread, with contributions from Alphaproteobacteria, Anaerolineae, Bacteroidia, Terriglobia, and Ignavibacteria, indicating active sulfur cycling that may regulate redox conditions and energy flow in the metabolic matrix.

**Fig. 2:**
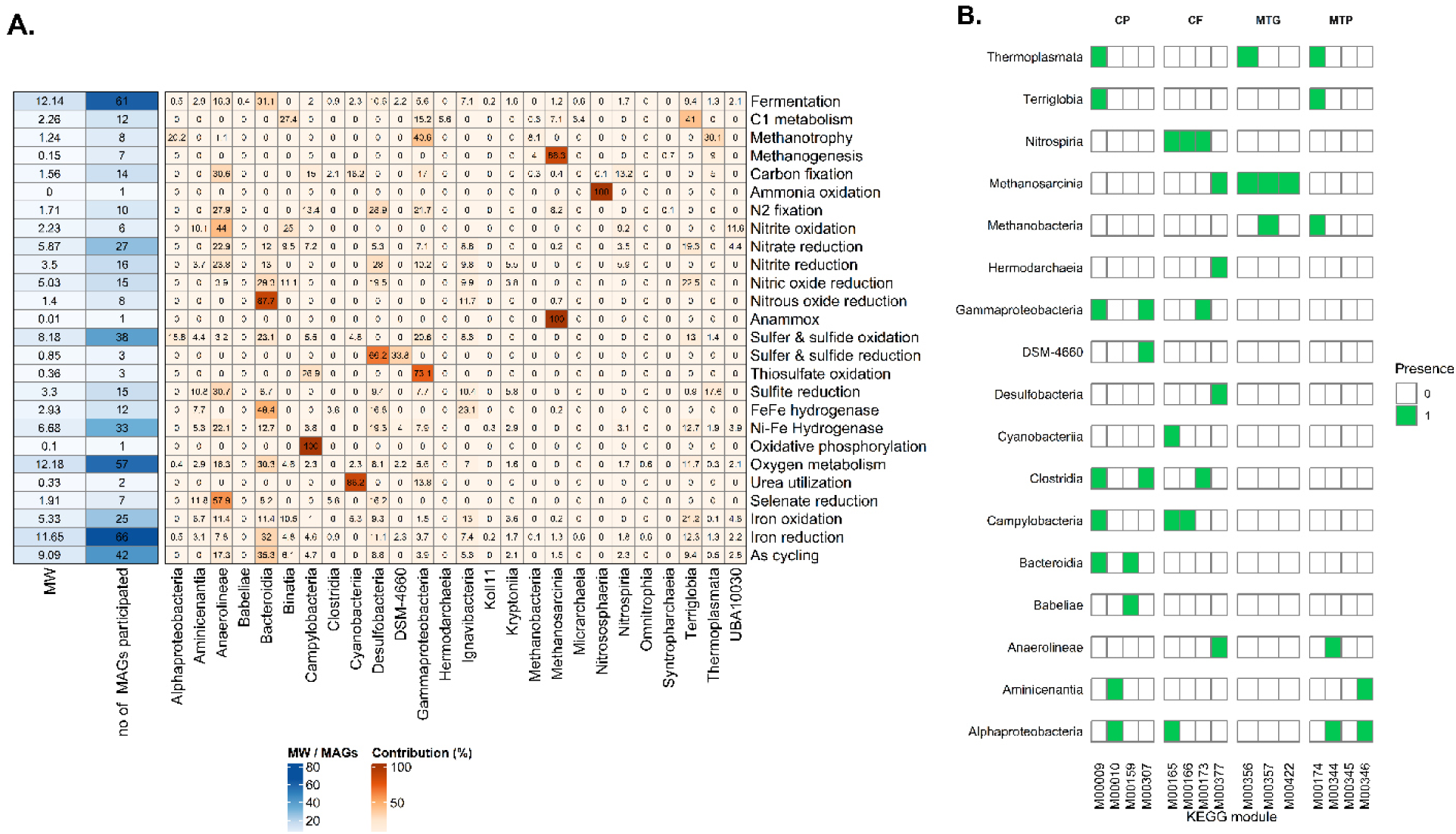
Metabolic profile and KEGG module distribution across high- and medium-quality metagenome-assembled genomes. Heatmaps showing metabolic weight score (MW) scores for each function, with the number of MAGs involved, along with the percentage contribution of each class for each function. Values in each cell represent the quantitative score or percentage contribution, with color gradients reflecting magnitude (A). Presence-absence of selected KEGG modules across MAGs, grouped by key functional categories (CP: CO₂ production, CF: CO₂ fixation, MTG: Methanogenesis, MTP: Methanotrophy). Each tile indicates whether a microbial class harbors the module completely (green) or not (white) (B).

To further examine the distribution of CO₂ and CH₄ cycling pathways at the module level, we considered KEGG module step hits obtained from METABOLIC v4. Among the KEGG modules, M00009 (TCA cycle), M00377 (Wood-Ljungdahl pathway), M00357 (acetoclastic methanogenesis), and M00174 (aerobic methane oxidation: methane => formaldehyde) were dominant across MAGs for CO₂ production, CO₂ fixation, methanogenesis, and methanotrophy, respectively (Fig. 2B). Among CO₂ production modules, two complete modules were present in Gammaproteobacteria, Clostridia, and Bacteroidia, whereas Nitrospiria encoded three complete modules among four CO₂ fixation paths. In contrast, all complete methanogenesis modules were restricted to Methanosarcina, with Methanobacteria and Thermoplasmata encoding only a single module corresponding to the methylotrophic and hydrogenotrophic pathways, respectively. For methanotrophy, Alphaproteobacteria contained two complete modules, while other taxa encoded a single module each.

### Temporal changes in CO_2_ and CH_4_ metabolic potential in prawn ponds

Based on KEGG-annotated gene functions detected across all samples, we reconstructed an integrated community-level metabolic network representing the genetic potential for CO₂ and CH₄ cycling in prawn pond ecosystems, linking central carbon metabolism, C₁ transformations, and methane turnover (Fig. 3). CO₂ production potential was primarily driven by oxidative reactions centered on pyruvate and acetyl-CoA metabolism, including malate and oxaloacetate decarboxylation (*maeAB*, *oadAB*) and pyruvate oxidation via pyruvate:ferredoxin oxidoreductase (*porA,B,C,D,G*). Additional genetic potential for CO₂ release was indicated by the presence of acetate-forming pathways and formate oxidation (*fdhB*). In parallel, genes encoding multiple CO₂ fixation pathways were detected, including the Calvin–Benson–Bassham cycle (rbcL/S), the reductive acetyl-CoA (Wood–Ljungdahl) pathway (cdhA/cooS, acsB), and acetyl-CoA carboxylation (accA–D). Methane cycling was represented at the genetic level by both production and oxidation pathways, forming an internal carbon-recycling loop. Methanogenic potential encompassed hydrogenotrophic, acetoclastic, and methylotrophic routes, with CO₂ reduction to methane through canonical archaeal pathways involving fwd/fmd, *mtrA–H,* and methyl-coenzyme M reductase (*mcrA,B,G*) genes. Conversely, methane oxidation potential was supported by the detection of particulate and soluble methane monooxygenases (*pmoA,B,C* and *mmoB,C,D,X,Y,Z*), followed by genes involved in methanol and formaldehyde oxidation (*mxaF,L/mdh, fae*), linking CH₄ turnover to downstream CO₂-producing pathways at the genomic level.

**Fig. 3:**
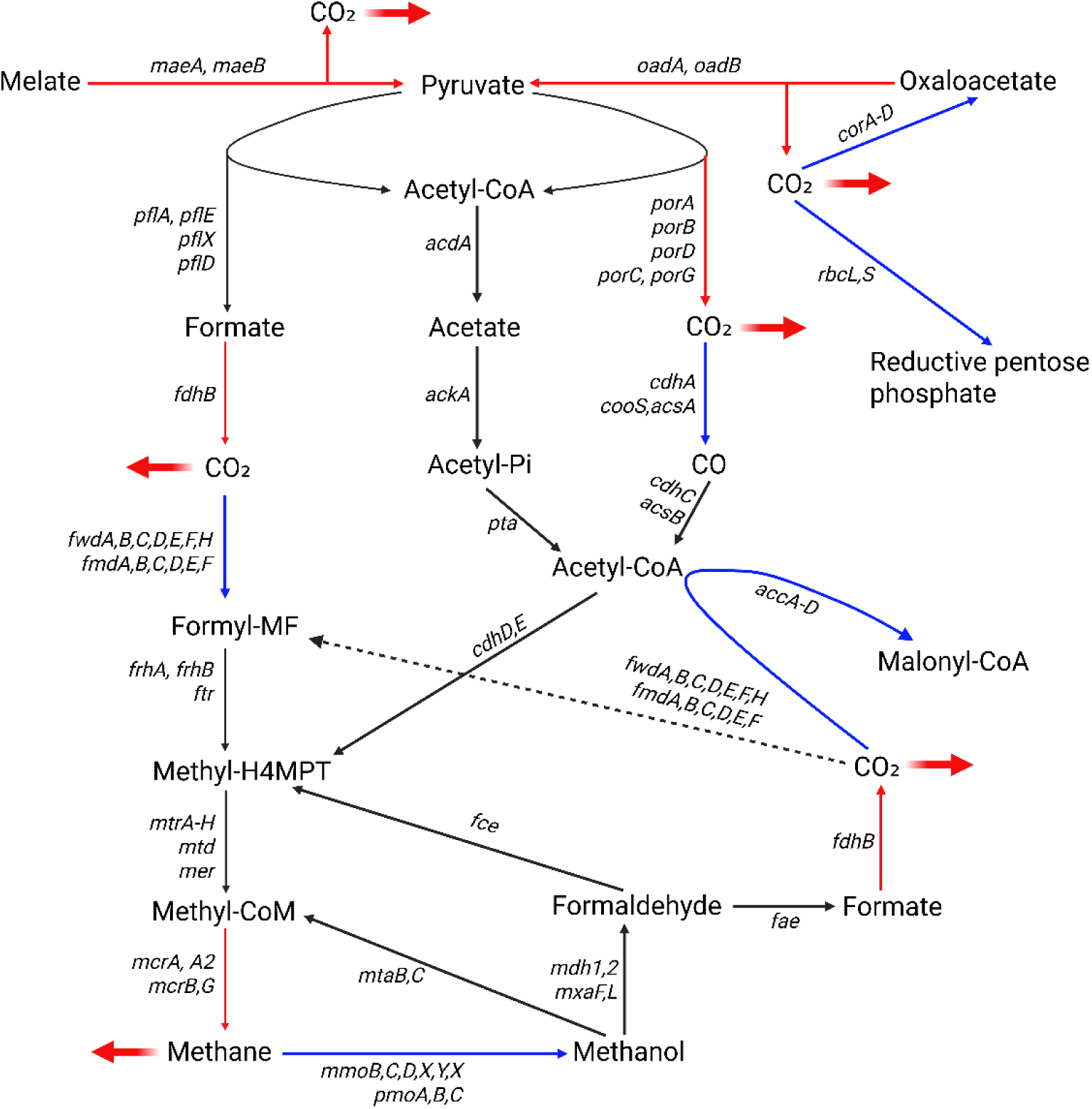
Integrated CO₂ and CH₄ metabolic potential network in prawn pond metagenomes. Key metabolic pathways and associated genes are shown based on gene evidence from recovered MAGs across all samples. Green and red arrows indicate the terminal steps of CO₂/CH₄ production and fixation/oxidation, respectively.

To discern seasonal- and treatment-driven shifts in metabolic potential, we quantified the abundances of key genes catalyzing the terminal steps of four major pathways: CO₂ production, CO₂ fixation, CH₄ production, and CH₄ oxidation. Clustering of terminal-step gene abundances revealed strong seasonal effects on functional gene profiles (PERMANOVA, R² = 0.64, F₅,₃₀ = 10.49, p < 0.001). K-means clustering of functional gene profiles revealed that early seasons (S1–S3) consistently grouped together, mid-late seasons (S4–S5) showed intermediate profiles, and the late season (S6) formed a distinct cluster across all four metabolic functions (Figure 4A and Supplementary note 4). In contrast, treatment had no significant effect on functional gene composition (R² = 0.06, F₅,₃₀ = 0.39, p = 0.94). Gene abundances increased progressively across seasons, peaking in S6, with significant differences between early (S1–S2) and late (S5–S6) seasons (Fig. 4B). Treatment-wise comparisons were non-significant, and differences among treatments were limited to modest shifts in mean gene abundances without consistent patterns across pathways (Fig. 4C). At the gene-specific level, *fdhB* and *cdhA* were the most abundant genes associated with CO₂ production and fixation potentials, respectively (see Supplementary note 5 for gene-specific trends), indicating that CO₂ production via formate oxidation and fixation through the Wood–Ljungdahl (W–L) pathway represent prominent genetic capacities within prawn pond microbial communities. In contrast, among CH_4_ cycling genes, *mcrA2* (non-canonical methanogenesis) and *pmoB* (methane oxidation) showed higher abundance compared to other genes in their respective groups. These genes showed the highest abundance in the late season (S6), with CO₂-related genes consistently dominating the overall functional gene pool across seasons and treatments.

**Fig. 4:**
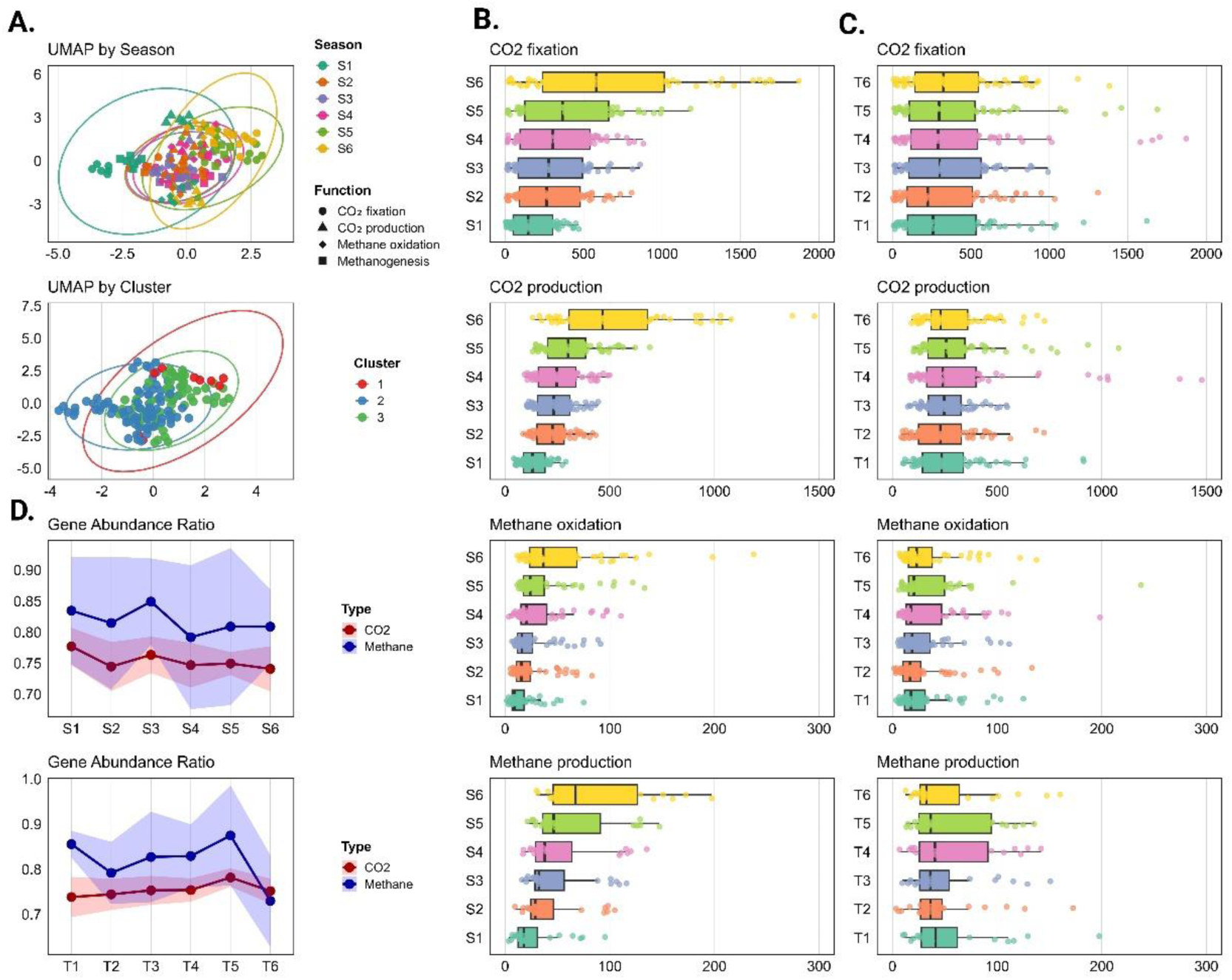
UMAP clustering, gene abundance, and functional ratios of CO_2_ and CH_4_ cycling genes in prawn pond metagenomes. UMAP projections display sample distributions based on CO₂ fixation, CO₂ production, methanogenesis, and methane oxidation gene abundances, colored by season and k-means cluster (A). Boxplots show gene abundances across seasons (B) and treatments (C), with color gradients denote grouping variables (seasons/treatments) and dots representing mean abundance per individual sample. Line plots illustrate mean gene abundance ratios (CO₂ production/fixation and methane production/oxidation) across seasons and treatments, with shaded ribbons indicating ± standard deviation (D).

To assess the trade-off between production and consumption potentials, we calculated abundance ratios of production-to-fixation/oxidation genes. For CO₂, the ratio was highest in S1 (0.78) and lowest in S6 (0.74), indicating a progressive increase in fixation potentials relative to production across seasons (Fig. 4D). The ratio for CH₄-related genes peaked in S3 (0.85) before declining. Across treatments, the largest trade-offs for both CO₂ and CH₄ related genes occurred in T5 (0.78 and 0.88, respectively) and T6 exhibited the lowest ratio for CH_4_ related genes (0.73), suggesting enhanced methane oxidation potentials in IMTA treatment (Fig. 4D).

### Microbial assemblages harboring CO₂ and CH₄ cycling genes respond to seasonal changes

To resolve the taxonomic origins of key CO₂ and CH₄ cycling genes, all genes were mapped to MAGs, revealing Bathyarchaeia as the most functionally versatile taxon, encoding 13 genes spanning CO₂ fixation and production, followed by Bacteroidia (10 genes), Gammaproteobacteria (8 genes), and Alphaproteobacteria (6 genes) (Fig. 5). Functions of all target genes are provided in Extended data ST3. CO₂ fixation genes, including *cdhA, cooS/acsA, fdhB, fwdB,E,F, fmdB,E,F, korA-d*, and *rbcL*, were primarily affiliated with Bathyarchaeia, Bacteroidia, Clostridia, Campylobacteria, and Thermoplasmata, whereas minor contributions came from Lokiarchaeia, Methanomicrobia, Methanosarcinia, Spirochaetia, and Anaerolineae. CO₂ producing *maeA,B, oadA, pdhA,B, porB-D,G,* genes originated mainly from Bathyarchaeia, Bacteroidia, Gammaproteobacteria, and Alphaproteobacteria, while methane producing genes, including *mcrA,B* weres exclusive to archaeal, dominated by Methanosarcina, Methanomicrobia, Thermoplasmata, and Methanomethylicia. In contrast, methane oxidation genes (*mmoB,C,D,Y,Z* and *pmoA,B,C*) were bacterial, represented mainly by Gammaproteobacteria and Alphaproteobacteria, with minor contributions from Methylomirabilia, Kiritimatiellia, Verrucomicrobia, Leptospiria, and Spirochaetia.

**Fig. 5:**
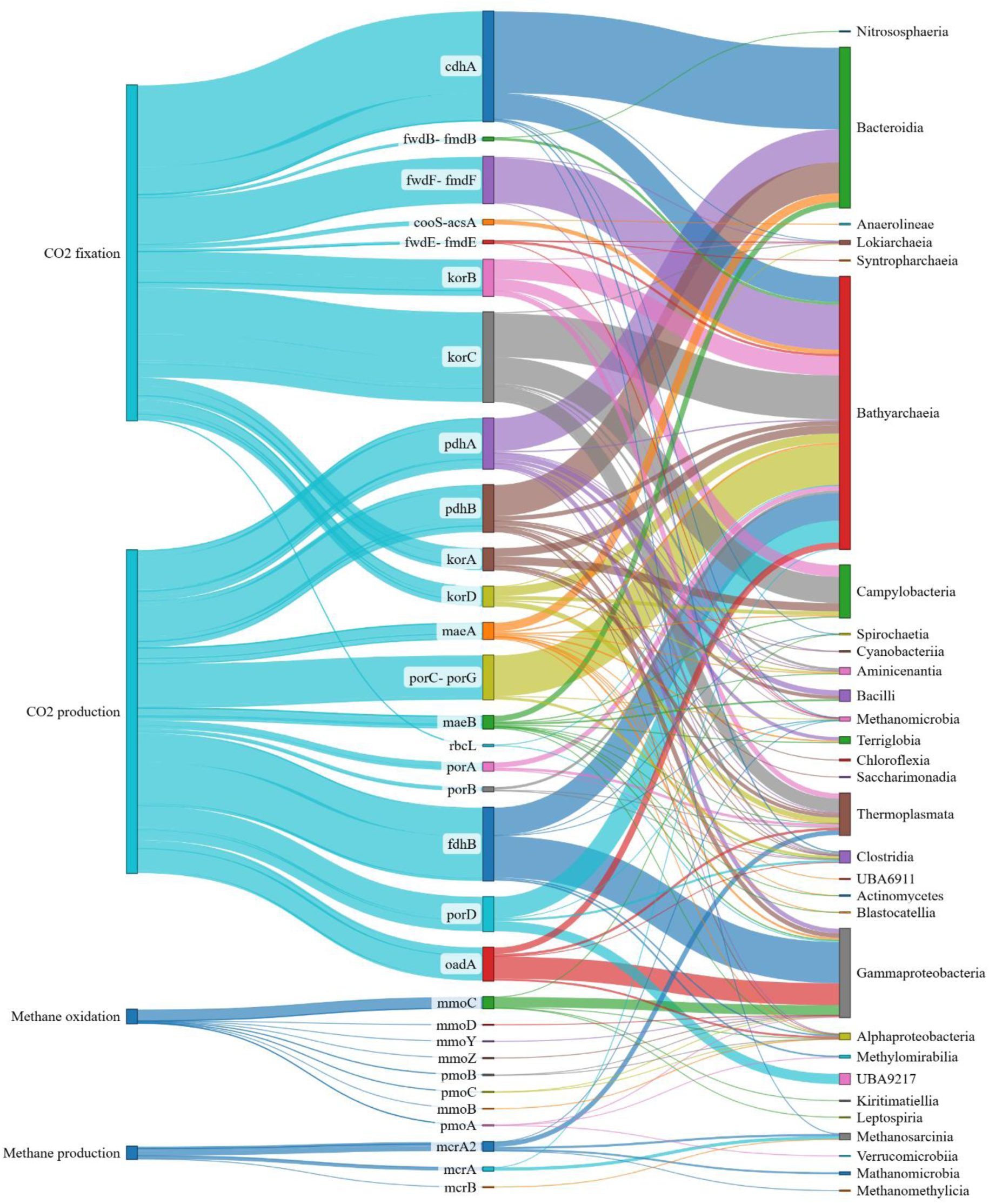
Sankey diagram showing the linkage between gene functions, individual genes, and microbial classes in prawn pond metagenomes. The diagram connects KEGG-annotated functions involved in CO₂ and CH₄ cycling to the corresponding genes, which are further linked to their microbial taxonomic classes. Line widths are proportional to the number of genes observed per function/class pair, providing a quantitative view of functional contributions across taxa.

With these core taxa, we examined how microbial community structure and composition varied across sampling occasions and treatments. Shannon diversity deviated significantly from normality (Shapiro–Wilk test, W = 0.85, P < 0.001) and showed heteroscedasticity across both seasons (Bartlett’s test, P = 0.004) and treatments (P = 0.015); therefore, non-parametric KW tests were applied. Shannon diversity differed significantly among seasons (P = 0.003), with S4–S6 exhibiting higher diversity than S1 but was unaffected by treatment (P = 0.48) (Fig. 6A). Bray–Curtis PERMANOVA confirmed that season structured microbial composition (adonis2, F₅,₃₀ = 3.51, R² = 0.37, P = 0.001), whereas treatment had no significant effect (F = 0.79, R² = 0.12, P = 0.84). Seasonal differences were accompanied by significant heterogeneity in community dispersion (PERMDISP, F = 2.74, P = 0.032) and NMDS ordination revealed a clear separation of late-season communities (S6) from early-season communities (S1–S2), with mid-season samples showing partial overlap (Fig. 6B).

**Fig. 6:**
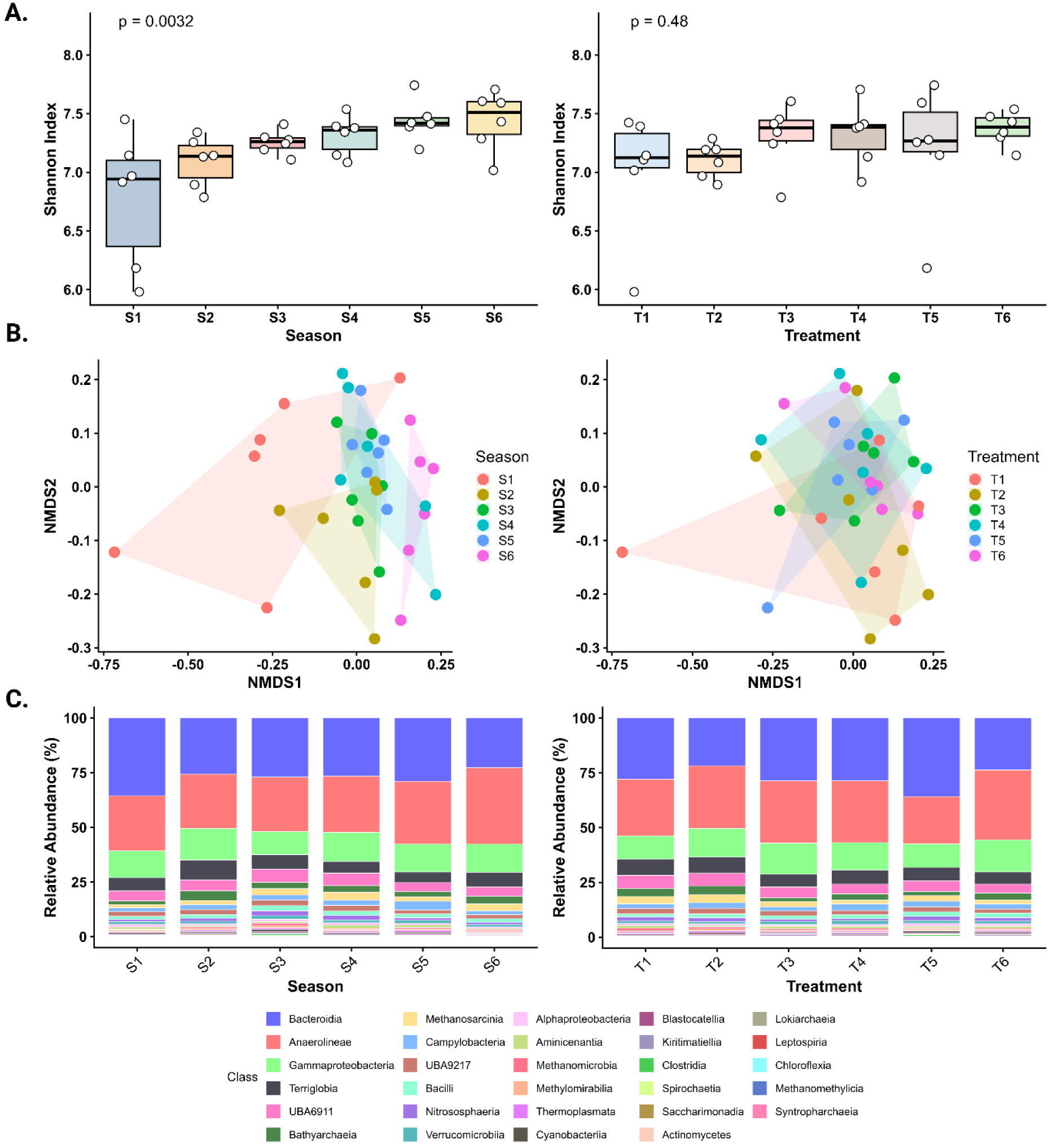
Microbial diversity, community structure, and relative abundance of MAGs across seasons and treatments. Boxplots show the Shannon index of MAGs by season and treatment (A), with medians and interquartile ranges; individual samples are overlaid as points. Statistical significance was assessed using KW. Non-metric multidimensional scaling (NMDS) based on Bray–Curtis dissimilarities of MAG abundances is shown in (B), with polygons indicating the convex hull of each season or treatment group; stress values indicate goodness of fit. Relative abundance of microbial classes across seasons and treatments is shown in (C). Data were normalized to percent abundance per sample, and colors correspond to microbial classes, ordered by total abundance across all samples (see more information in Supplementary note 6).

The microbial community was dominated by Anaerolineae (25–35%), Bacteroidia (23–36%), and Gammaproteobacteria (10–15%) across all samples, consistent with the dominant microbial classes represented among the reconstructed high-quality MAGs. Genome-resolved analyses further showed that Anaerolineae accounted for the largest proportion of H-MQ MAG abundance (19.99%), followed by Terriglobia (14.42%), Desulfobacteria (10.79%), Ignavibacteria (8.64%), Gammaproteobacteria (6.87%), and Bacteroidia (5.38%). Although Terriglobia and Desulfobacteria were among the dominant MAG classes, they contributed little to the targeted carbon-cycling pathways because few taxa belonging to Terriglobia encode genes associated with carbon-cycling processes, whereas Desulfobacteria lacked direct genetic potential for carbon-cycling pathways. Consequently, these taxa were abundant members of the microbial community but were not dominant contributors to the targeted carbon-cycling functions. In contrast, Syntropharchaeia exhibited consistently low relative abundance in both the reconstructed MAGs and the functionally annotated carbon-cycling taxa.Minor taxa responded primarily to temporal cues, with Alphaproteobacteria (0.65–1.30%), Syntropharchaeia (0.00015–0.00135%), and Lokiarchaeia (0.05–0.11%) showing significant seasonal shifts, while Methanosarcinia (1.7–3.6%) and Actinomycetes (0.05–0.17%) exhibited marginal changes (Fig. 6C). These temporal patterns were broadly consistent with the genome-resolved analysis, where Alphaproteobacteria and Methanosarcina MAGs also increased significantly across seasons. Experimental manipulation minimally changed class-level relative abundances (all P > 0.13), although Gammaproteobacteria and Methanosarcina were relatively higher and lower, respectively, in snail-containing treatments (T3 and T6). At the class level, most taxa increased progressively across sampling occasions, while Clostridia, Cyanobacteria, and Syntropharchaeia were relatively more enriched in monoculture treatment (T5) (Supplementary note 6).

## DISCUSSION

At the ecosystem level, GHG flux measurements from the same experimental system revealed pronounced temporal and treatment-level variability (Supplementary Note 1 and Ref^9^). CO₂ fluxes showed a general increasing trend over the production cycle, coinciding with progressive biomass accumulation and organic matter loading, whereas non-aquaculture control ponds maintained comparatively lower and more stable emission rates. In contrast, CH₄ fluxes increased during the early-to-mid production period (up to S4) and declined toward the end of the cycle, consistent with seasonal shifts in temperature and precipitation. Although differences among culture treatments were observed, most notably higher CO₂ and CH₄ fluxes in T1 and T5, respectively.

While bacteria and archaea are widely recognized as central microbial regulators of CO₂ and CH₄ cycling and emissions in aquaculture ponds, their community structure, metabolic pathways, variation in functional potential across production cycles, and response to culture management remain unexplored in prawn farming systems. As the first metagenomic investigation of prawn ponds, our study provides a comprehensive characterization of pond microbiomes, linking taxonomic composition to functional potential for CO₂ and CH₄ cycling across both temporal and management gradients. The recovery of 78 HMQ MAGs (63 bacterial and 15 archaeal) spanning diverse microbial lineages underscores the complexity of pond microbiomes and reveals previously underappreciated taxa, including Bathyarchaeia and Terriglobia, as potential mediators of carbon transformations in aquaculture environments. This enhanced resolution likely reflects our integrative metagenomic approach across the entire production cycle, an aspect not captured in earlier amplicon and snapshot metagenomic surveys^7,31,54^. The predominance of Bacteroidia, Anaerolineae, and Gammaproteobacteria among bacterial MAGs, together with Methanosarcina and Thermoplasmata among archaeal MAGs, indicates a microbial consortium structured by coexisting aerobic and anaerobic niches, consistent with observations from other aquaculture and sedimentary ecosystems^36,55–58^. Temporal variation strongly influenced MAG abundances, likely driven by progressive accumulation of organic and inorganic substrates in pond sediments. Remarkably, sulfate-reducing Desulfobacteria and DSM lineages increased significantly in relative abundance toward the late production season (S6) and were comparatively lower in non-feeding treatments (T2), suggesting possible stimulation by sulfur-rich organic inputs derived from feeds. These taxa likely promote localized anaerobiosis, with cascading effects on sulfur cycling, methanogenesis, and methanotrophy, thereby shaping CH_4_ dynamics within ponds^21–23^. In contrast, Gammaproteobacterial MAGs were relatively more abundant in lower organic environments, including IMTA ponds where organic matter was partially recycled by organic extractive species such as snails.

Our results demonstrate that CO₂ and CH₄ cycling pathways constitute only a minor fraction of the overall metabolic potential of prawn pond microbiomes, with methanotrophy, methanogenesis, carbon fixation, and C₁ metabolism collectively contributing far less than dominant heterotrophic processes such as fermentation, oxygen metabolism, and iron reduction. This carbon cycling metabolic weight score is lower compared to permafrost ecosystem^25^, wetlands^59^, peatlands^61^, deep ocean^60^, and human gut microbiome^61^. However, this metabolic structure is consistent with observations from other organic-rich aquatic systems^28,36,51^ and suggests that prawn ponds function primarily as heterotroph-dominated environments, sustained by substantial organic inputs from feed and detritus. Under such conditions, genetic potential for carbon mineralization appears to outweigh that for autotrophic carbon assimilation and methane turnover. The high representation of MAGs encoding iron-reduction pathways further suggests abundant availability of terminal electron acceptors that may support anaerobic respiration and organic matter oxidation in pond sediment^62^. Such dominance of respiratory and fermentative pathways is consistent with highly managed, nutrient-rich permafrost^51,63^ and aquaculture environments^36^, in which microbial metabolism is oriented toward rapid degradation of organic substrates rather than net carbon sequestration. These dominant heterotrophic pathways generate intermediate metabolites that can serve as substrates for downstream CO₂ and CH₄ production^64^. For example, the prevalence of acetoclastic methanogenesis modules encoded by Methanosarcina suggests that acetate derived from fermentation is one of the primary substrates supporting methane production in prawn ponds. Although representatives of the archaeal class Thermoplasmata (including Thermoplasmatales) have been proposed to have lost methanogenic capacity^65–67^, our detection of methylotrophic methanogenesis modules (M00356+02; K00399, K00401, K00402) in Thermoplasmatales (Thermoplasmataceae family) suggests a possible genetic capacity for methane production within this lineage in aquaculture environments. However, because the presence of genomic modules does not necessarily imply functional activity, further validation using transcriptomic analyses, stable isotope approaches, or controlled incubation experiments will be required to confirm active methanogenesis in aquaculture environments. Consistent with other aquatic and deep-sea systems, we identified sulfate-reducing Desulfobacteria and DSM lineages as potential mediators of sulfate and sulfide reduction and as important biological sinks for dark carbons^68^. Owing to their broad enzymatic capabilities^68–70^, these taxa were involved in multiple metabolic processes, including carbon, nitrogen, and sulfur cycling. Under anoxic conditions, sulfate-reducing bacteria can utilize Calvin-cycle–related enzymes to assimilate available carbon substrates^70^, thereby potentially reducing CO₂ availability and indirectly constraining downstream methane production by methanogens. This highlights the crucial role of sulfur cycling microbes in regulating GHG dynamics in aquaculture ponds and the need to explicitly integrate it into future assessments of genetic potential for aquaculture-derived GHG emissions.

One of the central objectives of this study was to reconstruct an integrated CO₂ and CH₄ cycling metabolic network representing the potential GHG production and consumption pathways in prawn ponds. Our results suggest that CO₂ production potential is mediated by multiple oxidative reactions, highlighting the predominance of heterotrophic carbon oxidation. The widespread detection of genes involved in formate oxidation and acetate-producing pathways further indicates that fermentation-derived intermediates may represent important substrates contributing to CO₂ release, a pattern broadly consistent with intensive feed inputs and organic matter accumulation in managed aquaculture systems^71^. Among the six known microbial CO₂ fixation pathways, the coexistence of five pathways, including the Calvin–Benson–Bassham cycle, Wood–Ljungdahl pathway, reductive TCA cycle, 3-hydroxypropionate bicycle, and 4-hydroxybutyrate pathway, demonstrates substantial diversity and flexibility of metabolic potentials within prawn pond microbiomes. This apparent functional diversity likely reflects the strong heterogeneity of organic substrates and fluctuating redox conditions in pond sediments and water columns, enabling different microbial strategies for carbon assimilation to coexist^72^. Such functional redundancy aligns with more environmentally exposed, organic systems^16,73^ and suggests that prawn ponds support a highly adaptable microbial carbon network. Methane cycling genes collectively point to the potential for an internal carbon-recycling loop, with methanogenesis proceeding through hydrogenotrophic, acetoclastic, and methylotrophic routes, followed by rapid methane oxidation linking CH₄ back to CO₂. All methanogenic routes converged on the complete methyl-coenzyme M reductase complex (*mcrABG*) as the terminal step. Hydrogenotrophic methanogenesis relies on CO₂ reduction using H₂ as an electron donor, whereas acetoclastic methanogenesis depends on acetate conversion to acetyl-CoA. In line with previous findings^74^, we also noticed reverse *Ack/Pta* pathways encoded by certain bacteria, including *Geobacter*, *Chlorobi*, and *Bacteroidetes*, suggesting potential competition for acetate between bacterial heterotrophs and obligate acetoclastic methanogens such as *Methanothrix*. Genes associated with both *mta*- and *mtr*-dependent methylotrophic methanogenesis pathways were also present, consistent with prior reports identifying these routes as important components of methane production potential in aquatic sediments^75^. However, without transcriptomic evidence, the relative activity and dominance of these pathways cannot be discussed with certainty. On the other hand, evidence on methane oxidation was restricted to the genes related to RuMP pathway, encoded by type Ib (Gammaproteobacteria) and type II (Alphaproteobacteria) methanotrophs^76^, whereas key genes involved in the serine pathway were not present. Additionally, anaerobic methane oxidation potential was inferred from the presence of ANME-affiliated lineages, which are thought to operate methyl-coenzyme M reductase in reverse under anoxic conditions. These methane production and consumption potentials appear to be spatially structured along steep oxic–anoxic gradients within pond sediments and the water column. Although recent studies have challenged the paradigm that methanogens, particularly acetoclastic taxa, are strictly anaerobic^74,77,78^, we found no evidence of genes associated with aerobic acetate degradation (extradiol dioxygenase, *elh*; K03381) or oxidative-stress maintenance via rubredoxin (*rubr*; K03534) in archaeal MAGs.

Seasonal variation emerged as the dominant factor shaping CO₂ and CH₄ emission potentials, with strong clustering of terminal-step genes across production cycles and minimal treatment effects. The progressive increase in gene abundances toward the late season likely reflects cumulative organic matter loading, sedimentation, and redox stratification, which collectively can intensify both carbon mineralization and fixation^79^. Nevertheless, as these inferences are based on genomic potential rather than expression, we remain cautious in interpreting functional activity. Therefore, we also investigated production-to-consumption gene ratios, which provide a more robust gene-centric proxy for emission potential. The declining CO₂ production-to-fixation ratio across seasons suggests a gradual shift toward greater relative representation of carbon assimilation pathways, potentially associated with increasing availability of reduced carbon substrates, evolving redox gradients, and energy conditions favorable to chemoautotrophs^80^. Similarly, the mid-season peak followed by a decline in methane production-to-oxidation ratios broadly mirrors methane emission trends observed in the same experiment (Supplementary Note 1), although this correspondence should be interpreted cautiously.

The observed decline in ratios during the late season may reflect multiple, non-exclusive factors, including a relative increase in methanotroph-associated genes compared to methanogenic genes, reduced methanogenic potential under lower temperatures^81^ and elevated nitrogen levels^82^. An additional contributing factor could be the increasing abundance of sulfate-reducing and acetate-consuming bacteria over time, as observed in Fig. 1, which likely outcompete methanogens for substrates and energy, thereby constraining methanogen proliferation^83^. At the treatment level, the lowest methane production-to-oxidation gene ratio observed in IMTA treatments underscores the potential of integrated management to reduce methane emission potentials, likely by moderating organic loading and complementing dissolved oxygen to favor methane oxidation.

Previously unrecognized Bathyarchaeia emerged as the most metabolically versatile lineage, contributing extensively to both CO₂ production and fixation and methane production potential, highlighting its possible role as a central hub in pond carbon metabolism. In line with previous reports^65–67^, we also found that Bathyarchaeia and Thermoplasmata harbor the *mcr* gene in pond ecosystems. Contrary to our initial hypothesis, microbial diversity and community composition were not structured primarily by treatment but rather by seasonal progression. The higher diversity and distinct community structure observed in late-season samples suggest increasing niche differentiation as ponds mature. Progressive organic enrichment likely drives stronger redox stratification with expanded anaerobic conditions^84^ that can significantly favour the relative increase of Anaerolineae lineage in pond sediments that harbor CO₂ fixation genes. At the same time, well-defined oxic–anoxic interfaces may support aerobic processes, including methane oxidation *pmoA*-containing Gammaproteobacteria. These partly explain the reduced production-to-oxidation/fixation ratio observed in later seasons. The higher relative abundance of Gammaproteobacteria and lower relative abundance of Methanosarcina in snail-containing treatments likely reflects reduced organic loading and enhanced bioturbation or grazing by snails, which could promote more oxic conditions and favor aerobic or facultative heterotrophs and methanotrophs over strict anaerobic methanogens^85^. In contrast, the enrichment of Clostridia, Cyanobacteria, and Syntropharchaeia in monoculture ponds (T5) suggests greater organic accumulation and the development of stronger anaerobic niches, highlighting the potential for elevated CO₂ emissions in monoculture-based production systems.

Despite providing genome-resolved insights of microbial metabolic potential in prawn ponds, this study is limited by its reliance on metagenomic inference rather than transcriptomic or proteomic evidence, which prevents definitive conclusions about in situ metabolic activity and pathway dominance. Second, although we recovered a set of dereplicated high-quality MAGs spanning diverse bacterial and archaeal lineages, these genomes do not capture the full taxonomic and functional diversity present in these complex pond ecosystems. Even with these limitations, our findings demonstrate that CO₂ and CH₄ cycling potentials in prawn ponds are emergent properties of seasonally evolving microbial networks shaped by organic loading, redox stratification, and microbial competition, rather than the activity of individual taxa or pathways alone. Future studies integrating multi-omics approaches and improved genome recovery will be essential to resolve active processes and to elucidate functional synergies among sulfur–carbon–methane cycling microbial communities.

## Data availability

All data generated and/or analysed during the experiment are publicly available in the results sections and supplementary material.

## Authors contributions

AB, AMD, and LHH contributed to the study design and methodology. AB and MMH conducted field experiments and sampling. AB, AMD, and PDB performed lab activities, including library preparation for sequencing and bioinformatics. AB prepared figures and wrote the manuscript. MMH, LHH, and NOGJ managed funding, administered the project, and supervised the experiment. NOGJ, MHH, PDB, and MMRJ reviewed and edited the manuscript. All authors approved the submission of the manuscript.

