## Supplementary file for "Genome-resolved metagenomics reveals microbial potential for CO_2_ and CH_4_ emissions in prawn ponds"

**Supplementary Note-1: Water quality and GHG emission patterns in experimental ponds**

Seasons S2–S4 were characterized by heavy rainfall, whereas S5–S6 were dry with declining temperatures. Sustained rainfall and limited sunlight during mid-cycle redistributed organic matter and lowered sediment redox potential^1^, leading to declining DO until S4, but recovered later, with T4 and T6 maintaining higher DO levels throughout the production cycle. Water temperature followed the seasonal pattern, decreasing from the rainy to the dry periods. In consistent with previous studies^2,3^, TDS and TAN increased progressively with water age, with TDS consistently higher in aquaculture treatments than in the non-aquaculture control (Fig. S1).

Across the production cycle, CH4 and CO2 emission dynamics were jointly influenced by seasonal precipitation, water quality, and treatment-specific management. Lower feeding rates and reduced organic matter deposition initially suppressed CO₂ and CH₄ emissions during the early culture phase (S1-S2), despite increasing biomass (Fig. S2). During mid-season (S3-S4), CO₂ and CH₄ fluxes increased in feed-intensive treatments (T1 and T5), reflecting enhanced organic matter redistribution and reduced redox potential. As rainfall declined after S4, DO recovered most strongly in plant-based treatments (T4 and T6), coinciding with reductions in CH₄ fluxes, indicating enhanced methane oxidation under more oxic conditions. In contrast, the non-aquaculture control (T2) experienced smaller rainfall-driven shifts in water quality and consistently lower emission rates.

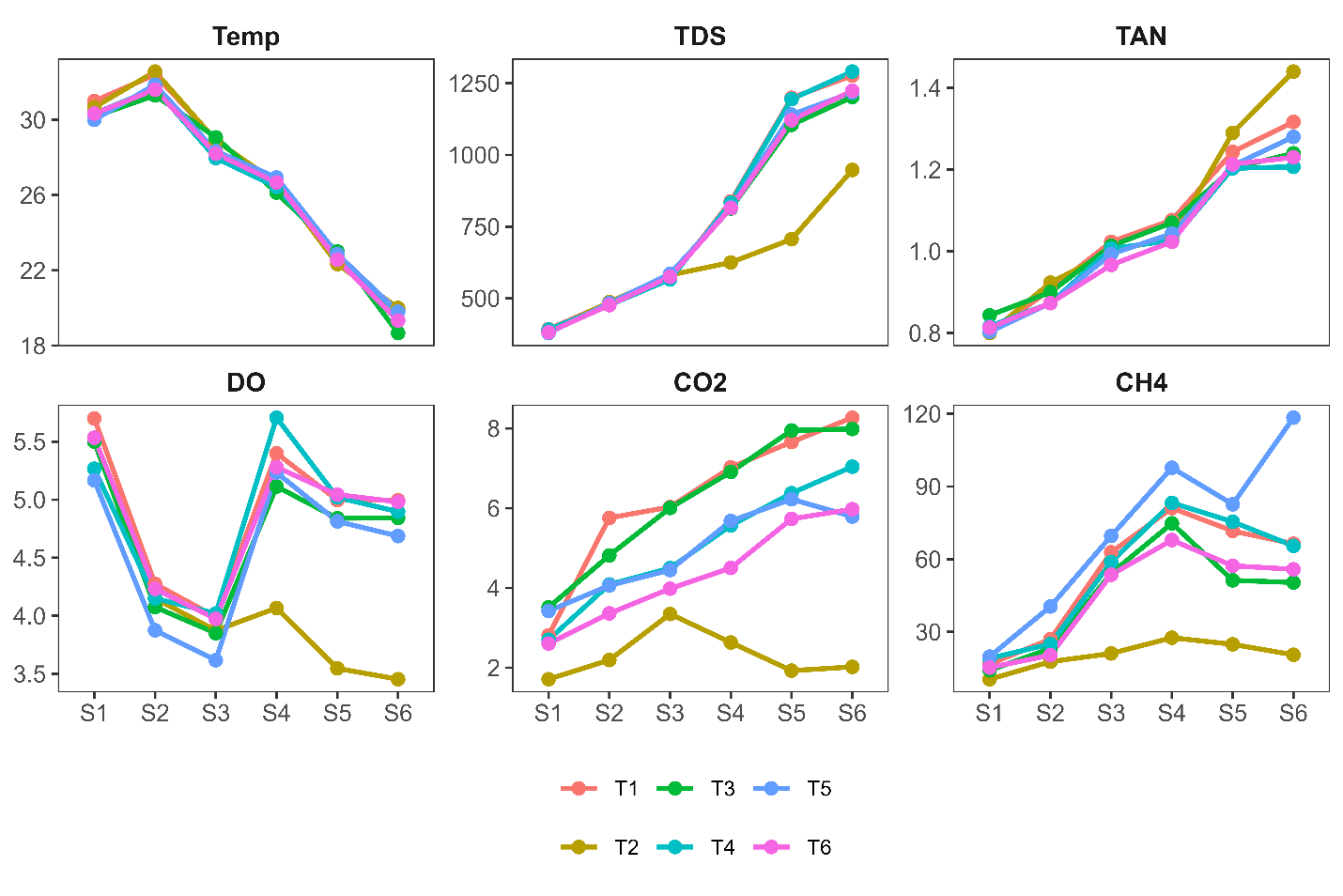

Fig. S1: Seasonal variation in water temperature (°C), total dissolved solids (TDS: ppm), total ammonia nitrogen (TAN: ppm), dissolved oxygen (DO: ppm), CO₂ fluxes (mg/m^2^/h), and CH₄ fluxes (μg/m^2^/h) across six seasons (S1–S6) for different pond treatments. Lines represent treatment-wise mean values across replicate ponds.

**Supplementary note 2: Adapted Hackflex Protocol for Shotgun Metagenomic Library Preparation**

Bead-linked transposomes (BLT; Illumina) were diluted 1:50 in molecular-grade water and stored at 4–8 °C. Tagmentation buffer was prepared in-house by combining 20 mM Tris-HCl, 20 mM MgCl₂, and 20% dimethylformamide in nuclease-free water, and stored at –20 °C. Tagmentation stop buffer (TSB) consisted of 0.2% SDS in water and was stored at room temperature. All reagents were equilibrated to room temperature before use as per the adapted Hackflex workflow. The workflow consisted of four main stages: tagmentation, stopping tagmentation, PCR amplification, and library cleanup (Fig. S2). For tagmentation, 5–10 ng of high-quality genomic DNA was added to PCR tubes and BLT beads and tagmentation buffer were added to prepare a Tagmentation Master Mix. The reaction was placed on a thermal cycler and incubated at 55 °C for 15 minutes with the lid preheated to 100 °C. The tagmentation reaction was halted using 10 µL of tagmentation stop buffer added to each tube. Beads were gently mixed 10 times using a pipette to ensure complete contact between the TSB and bead-bound transposomes. The tubes was incubated on a thermal cycler at 37 °C for 15 minutes, then held at 10 °C. Keeping on a magnetic stand, supernatant was discarded and PCR was performed directly on the tagmented DNA attached to BLT beads. PCRBio HiFi polymerase and custom dual-index i5 and i7 primers were used. Libraries were purified using Ampure XP beads and freshly prepared 80% ethanol. Beads were equilibrated to room temperature for at least 30 minutes before use. Cleanup involved a double-size selection to remove unincorporated primers, adapters, and small fragments. Following two washes with 180 µL ethanol, libraries were eluted in 22 µL nuclease-free water.

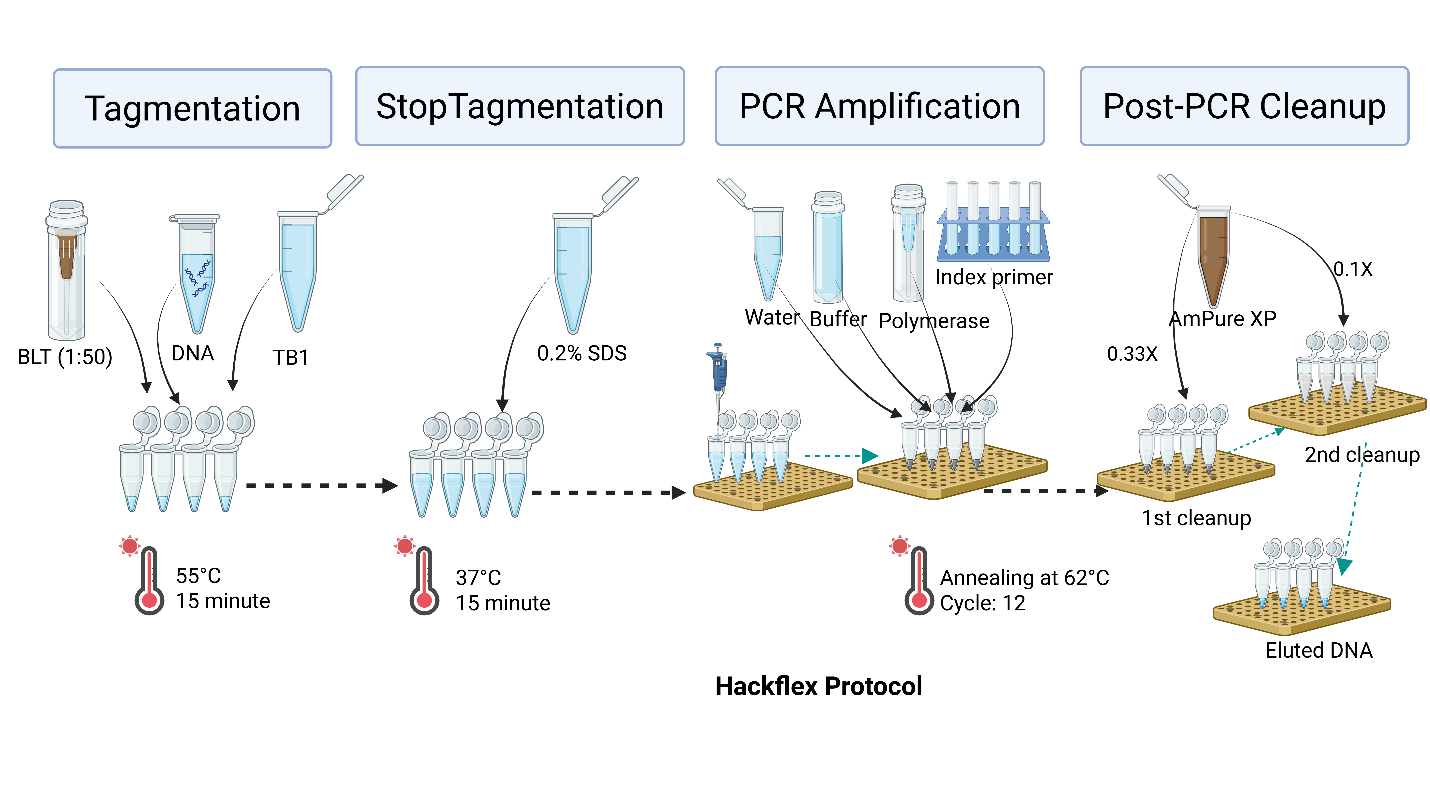

Fig. S2: Adapted heckflex protocol used in this study

**Supplementary note 3: Statistics on quality of recovered bins**

Metagenomic binning using the Taxvamb pipeline yielded a large collection of prokaryotic genome bins spanning a wide range of completeness and contamination levels. In total, 82,181 bins exhibited contamination levels ≤5%, whereas only a small fraction showed moderate (5–10%; n = 90), high (11–30%; n = 76), or very high contamination (>30%; n = 86), indicating generally effective binning performance (Table S1). Genome completeness estimates revealed that most bins were fragmented, with 81,348 bins falling within the 11–30% completeness range. In contrast, 152 bins exceeded 50% completeness, including 31 bins with completeness greater than 90%. Genome sizes were predominantly small, with the majority of bins less than 1 Mbp (n = 81,952), while fewer bins fell within intermediate (1–3 Mbp; n = 114) or larger size classes (3-5 Mbp; n = 41; >5 Mbp; n = 74), consistent with fragmented assemblies derived from complex environmental samples^4–7^.

Table S1: Quality and genome size of all recovered bins

|  | Range | No. of MAGs |
| --- | --- | --- |
| Completeness (%) | <10 | 408 |
|  | 11-30 | 81348 |
|  | 31-50 | 273 |
|  | 50-75 | 52 |
|  | 76-90 | 69 |
|  | >90 | 31 |
| Contamination (%) | <5 | 81929 |
|  | 5-10 | 90 |
|  | 11-30 | 76 |
|  | >30 | 86 |
| Genome size (Mbp) | <1 | 81952 |
|  | 1-3 | 114 |
|  | 3-5 | 41 |
|  | >5 | 73 |

Taxonomic classification across all recovered bins revealed substantial ambiguity, with 1,656 bins remaining unclassified at all taxonomic ranks. Among classified bins, assignments were most frequently resolved at higher taxonomic levels, including kingdom (n = 29,191) and phylum (n = 2,393), whereas resolution declined markedly toward lower ranks, with 26,749 bins lacking species-level assignment (Fig. S3).

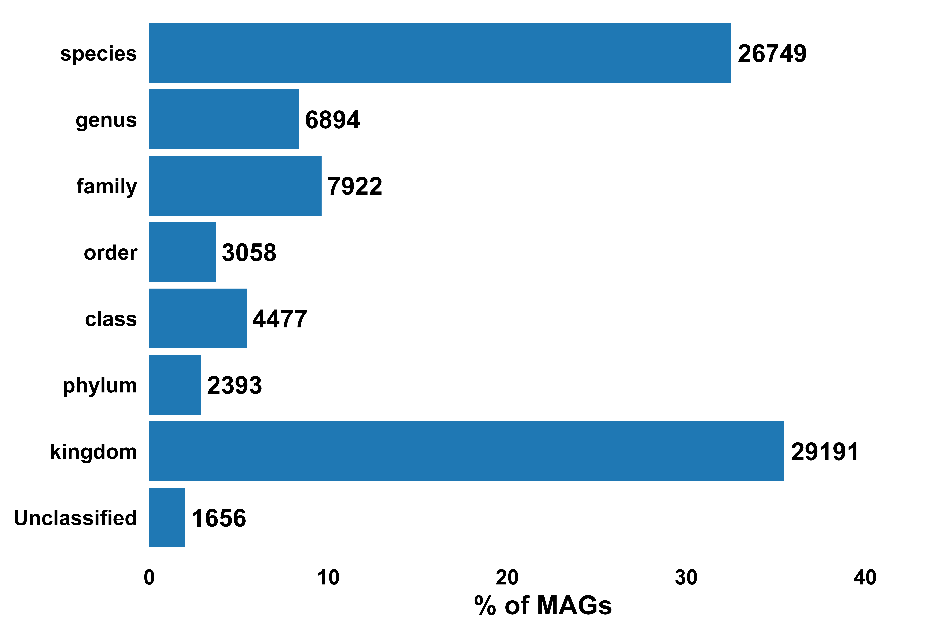

Fig. S3: The percentage of MAGs assigned to different taxonomic ranks, from domain to species level, based on the GTDB-TK2 database.

**Supplementary note 4: Clustering of gene abundance across seasons and treatments**

Clustering of CO₂ fixation and CO₂ production genes revealed clear seasonal structuring. For CO₂ fixation, Cluster 1 was exclusive to S6 (100%), indicating a distinct late-season profile. Cluster 2 included S5 (25%), S2 (20.8%), S3 (20.8%), S4 (20.8%), and S6 (12.5%), representing intermediate profiles, while Cluster 3 was dominated by S1 (66.7%), with minor contributions from S2, S3, S4 (11.1% each), forming a large homogeneous early-season group (Fig. S4). For CO₂ production, Cluster 1 was also exclusively S6 (100%), Cluster 2 included S4 (24%), S5 (24%), S2 (20%), S3 (20%), and S6 (12%), and Cluster 3 was mainly S1 (75%), with minor contributions from S2 and S3 (12.5% each). For methane oxidation, Cluster 1 contained only S6 (100%), Cluster 2 included S1–S3 (28.6% each) and S4 (14.3%), and Cluster 3 had S5 (46.2%), S6 (30.8%), and S4 (23.1%), indicating that late-season methane-oxidizing genes are more spread across clusters. Methanogenesis followed a similar trend: Cluster 1 contained only S6 (100%), Cluster 2 included S5 (45.5%), S6 (36.4%), and S4 (18.2%), while Cluster 3 comprised S1–S3 (26.1% each), S4 (17.4%), and S5 (4.35%).

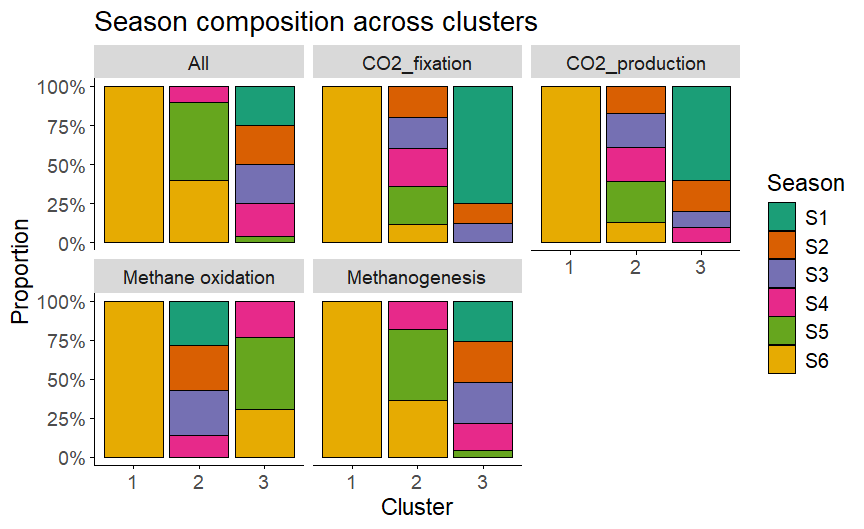

Fig S4: Hierarchical clustering of seasons based on the abundance of functional genes showing how different seasons influenced cluster composition for all four major functions.

**Supplementary note 5: Temporal and treatment-wise variation in gene abundances**

Season strongly influenced the abundance of functional genes involved in GHG emissions. Among the four functions analyzed, CO2 fixation (*cdhA*) and CO2 production (*korA*) genes were consistently the most abundant, followed by methane production (*mcrA2*) and methane oxidation (*pmoB*). Seasonal analysis revealed that S6 exhibited the highest gene abundances across all functions, with mean values of 1541 for *cdhA*, 1142 for *korA*, 171 for *pmoB*, and 234 for *mcrA2*. Post-hoc comparisons showed significant differences between early (S1–S2) and late seasons (S5–S6), particularly for CO2 fixation and production (p < 0.05), as well as for methane-related genes (p < 0.001), indicating enhanced microbial activity toward the end of the sampling period (Fig. S5).

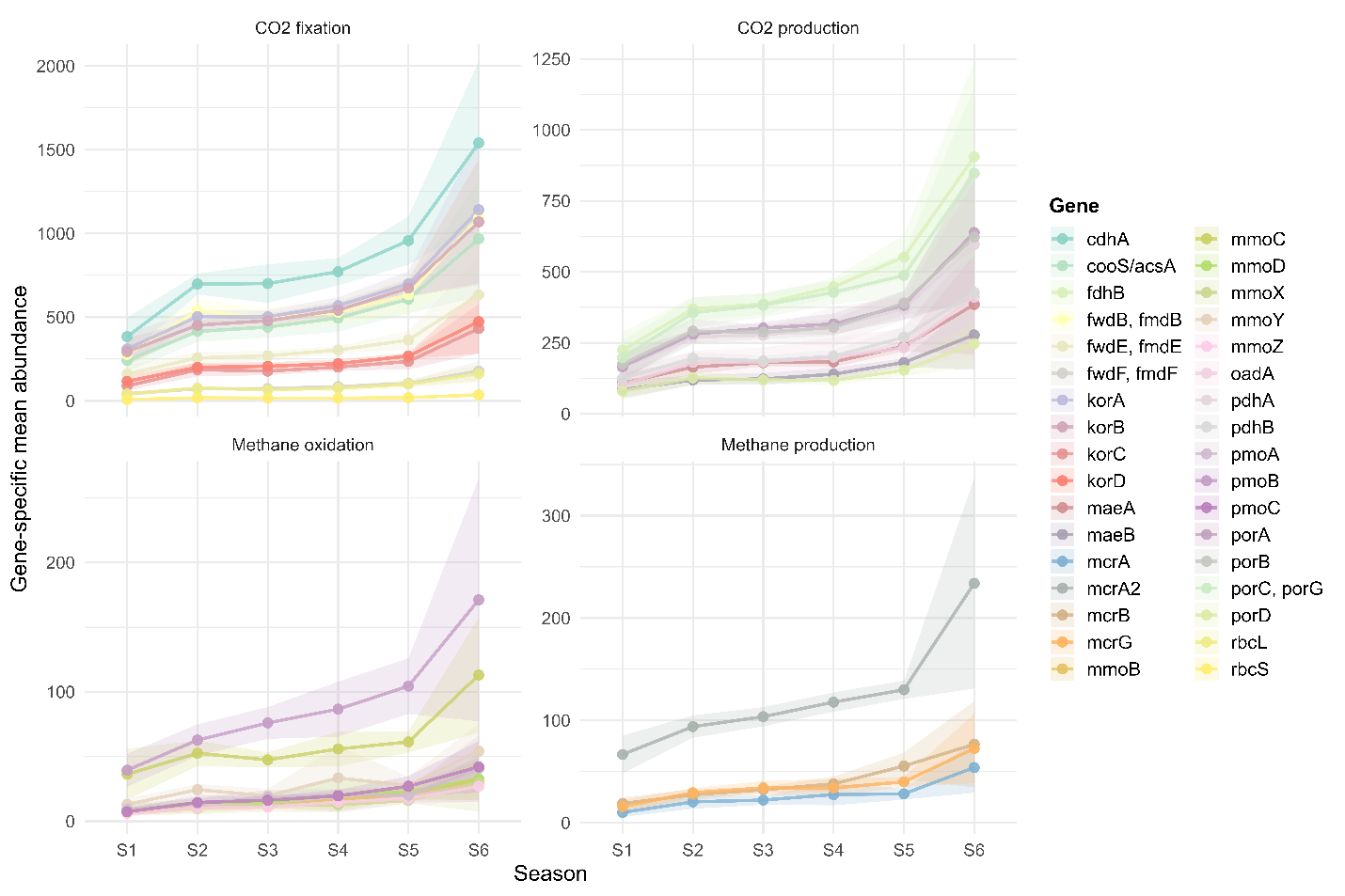

Fig. S5: Abundance of individual genes involved in key CO₂ and CH₄ cycling pathways across six seasons in prawn ponds.

Treatment effects were more moderate but still noticeable. T4 promoted the highest gene abundances for all functions, with cdhA at 996, korA at 738, pmoB at 121, and mcrA2 at 151. ANOVA results indicated that CO2 production was significantly affected by treatment (p = 0.024), whereas other functions showed no significant treatment effects. Pairwise post-hoc comparisons confirmed that T4 differed significantly from early treatments for CO2 production, but differences for other functions were not statistically significant (Fig, S6).

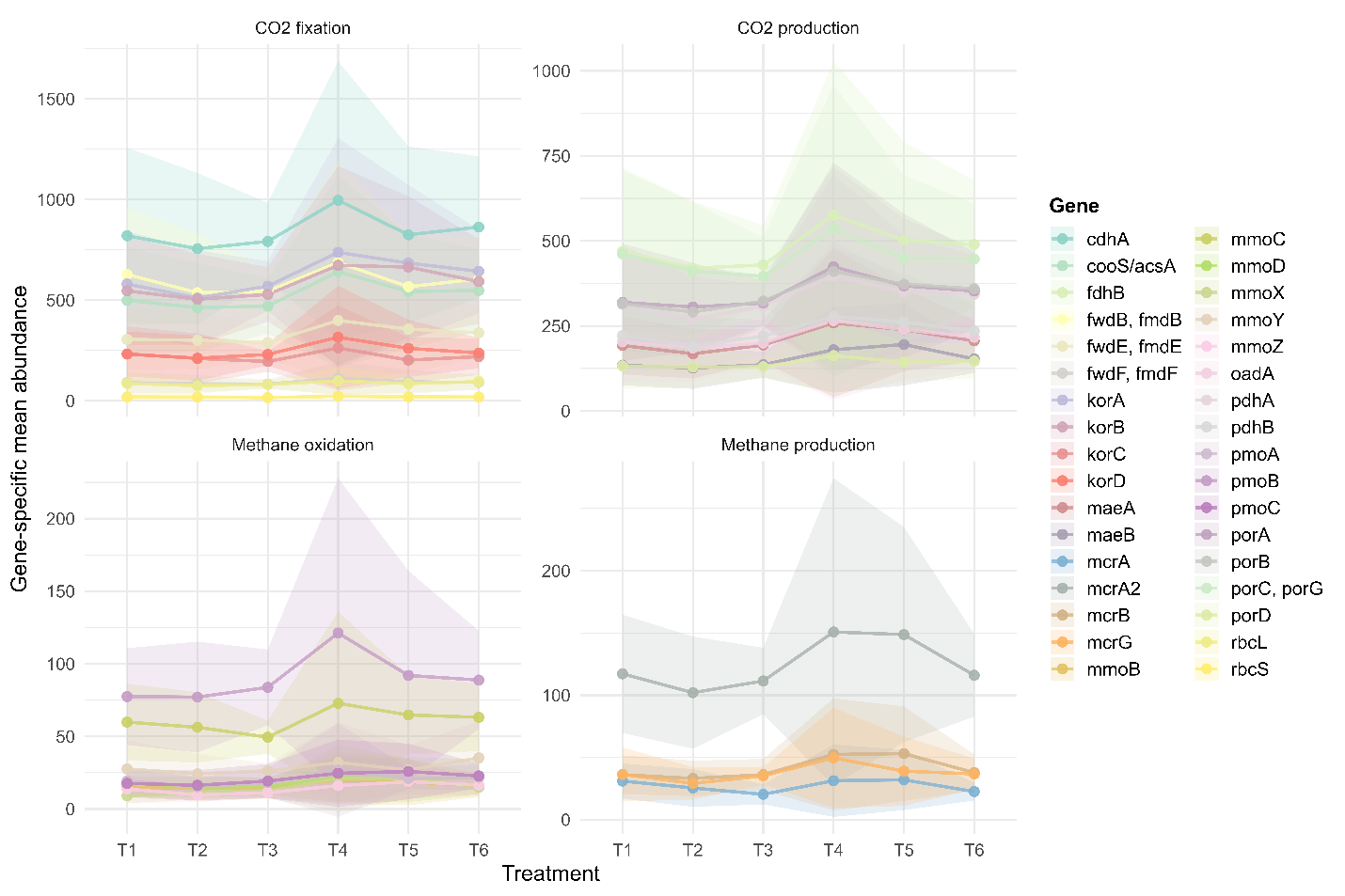

Fig. S6: Abundance of individual genes involved in key CO₂ and CH₄ cycling pathways across six treatments in prawn ponds.

Gene-specific trends showed that CO2-related genes (cdhA and korA) were generally more abundant than methane-related genes (pmoB and mcrA2) in both seasonal and treatment contexts. Line plots with standard deviation shading revealed that CO2 fixation and production increased sharply toward the late season and under T4 treatment, while methane oxidation and production showed a similar but less pronounced increase.

**Supplementary note 6: Relative abundance of core taxa across seasons and treatments**

Across all seasons, the community was consistently dominated by Bacteroidia and Anaerolineae, together accounting for 50–60% of total relative abundance, highlighting their central role in organic matter degradation under prevailing conditions. Bacteroidia were most abundant in early seasons (S1–S3) and declined toward S6, whereas Anaerolineae showed a clear increasing trend, reaching their maximum contribution in the late season (S6), indicating a seasonal shift toward more anaerobic, fermentative processes. Proteobacterial classes exhibited moderate but stable contributions across seasons, with Gammaproteobacteria (10–14%) remaining relatively constant and Alphaproteobacteria gradually declining from S1 to S6 (Fig. S7). Archaea classes associated with carbon cycling, including Bathyarchaea, Methanosarcina, Methanomicrobia, and Thermoplasmata, generally increased from early to mid–late seasons, suggesting enhanced anaerobic carbon transformation and methanogenic potential as the season progressed. Minor but ecologically relevant groups such as Nitrososphaeria, Methylomirabilia, and Campylobacteria peaked during mid- to late-seasons, indicating seasonal niche differentiation in nitrogen- and methane-related processes.

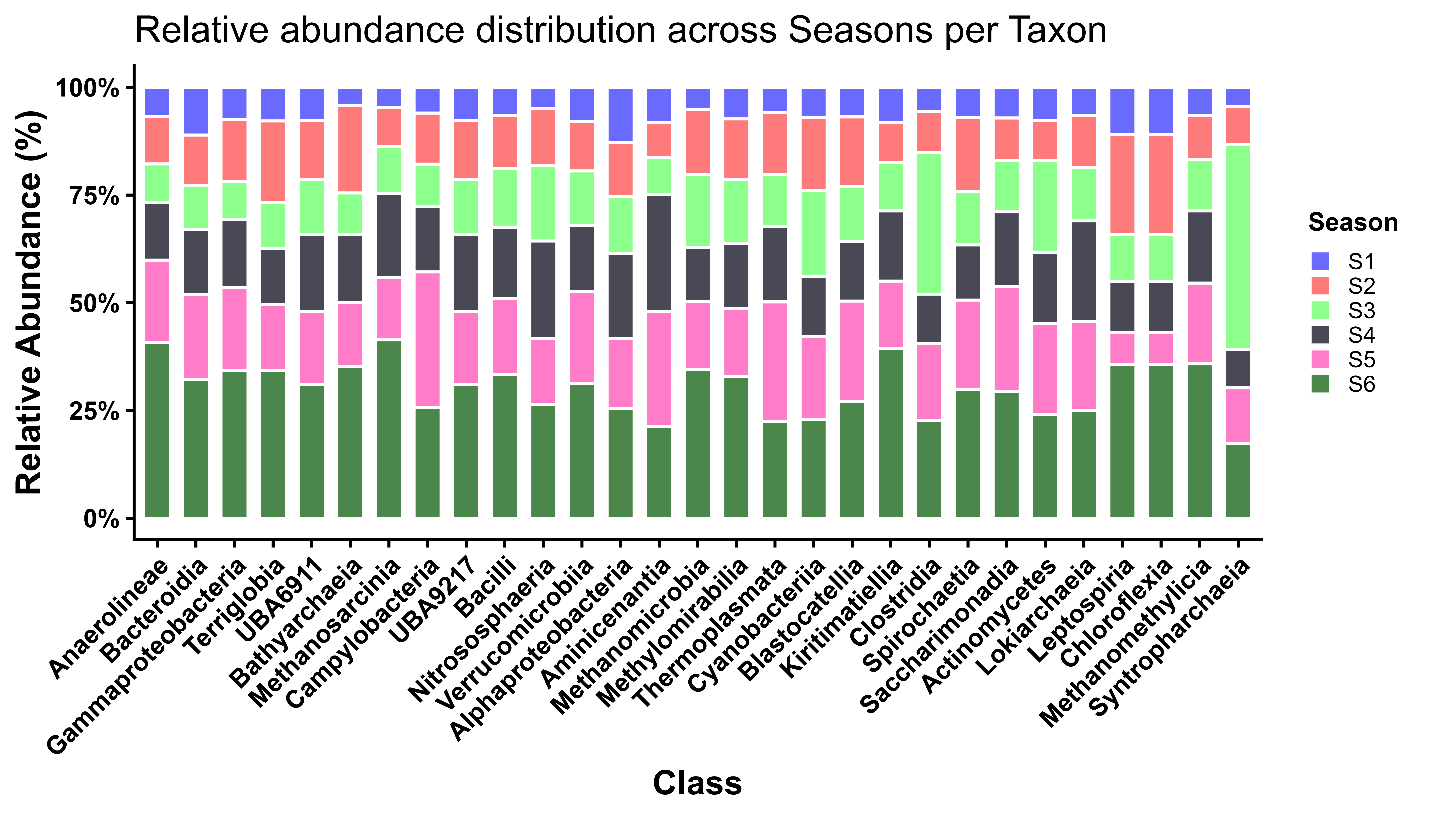

Fig. S7: Season-wise relative abundance of key microbial classes harboring genes responsible for the terminal steps of CO₂ and CH₄.

Across treatments, the microbial community retained its core structure, dominated by Bacteroidia and Anaerolineae, which together accounted for a substantial fraction of total abundance. Bacteroidia peaked in T5 and T4 but declined in T6, whereas Anaerolineae was highest in T6, highlighting treatment-specific shifts in fermentative taxa (Fig. S8). Archaeal classes involved in carbon cycling, including Methanosarcinia, Methanomicrobia, and Bathyarchaeia, showed variable responses across treatments, with higher relative abundance in T2–T3, suggesting modulation of methanogenic potential by treatment. Minor bacterial taxa such as Cyanobacteriia, Clostridia, and Campylobacteria also exhibited treatment-dependent fluctuations.

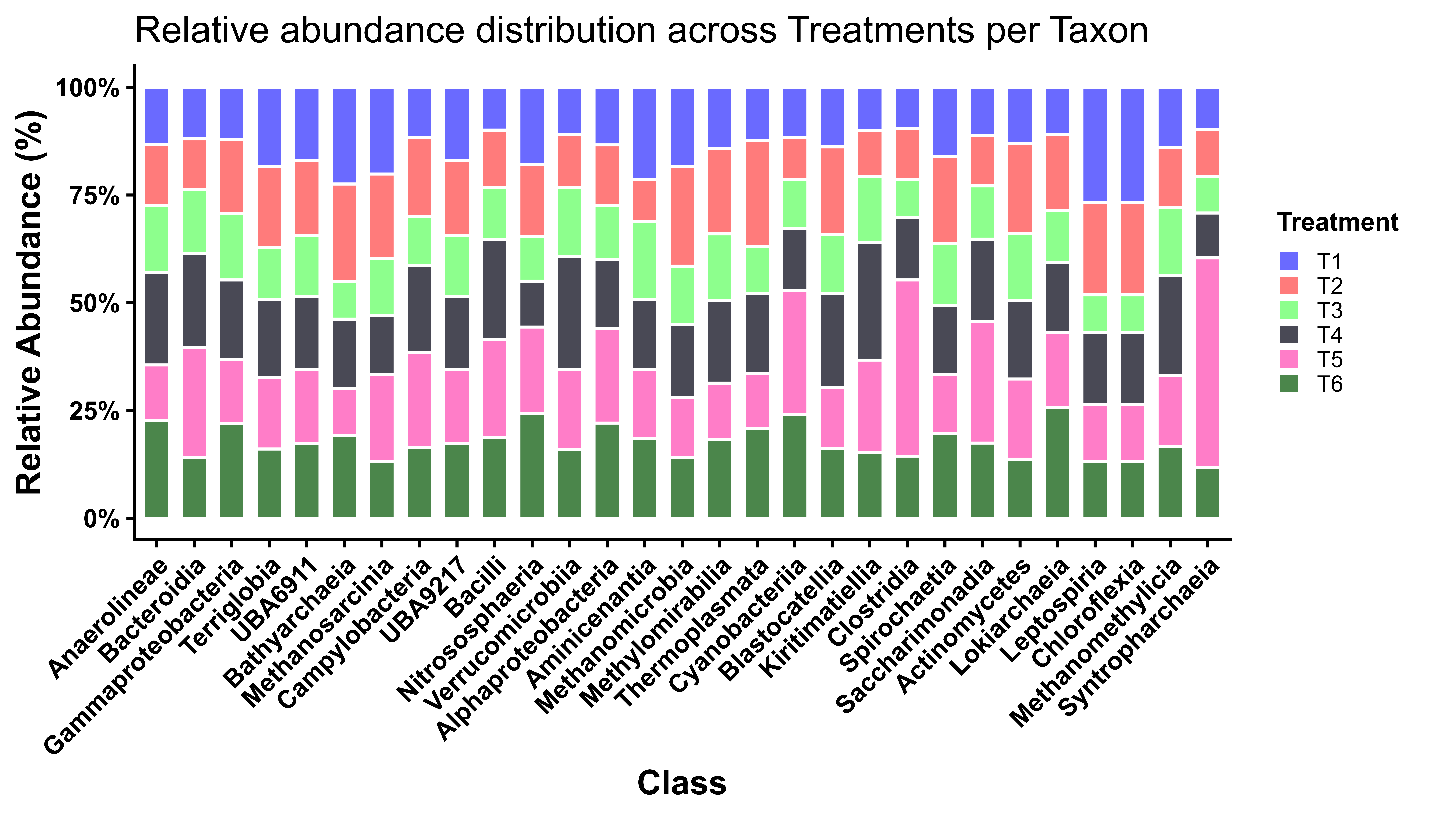

Fig. S8: Treatment-wise relative abundance of key microbial classes harboring genes responsible for the terminal steps of CO₂ and CH₄.
